# Evexomostat (SDX-7320), a Methionine Aminopeptidase Type 2 (METAP2) Inhibitor, Stimulates Weight Loss and Inhibits Obesity-Accelerated Tumor Growth

**DOI:** 10.64898/2026.01.28.702362

**Authors:** Peter Cornelius, Benjamin A. Mayes, Andrew J. Dannenberg, Pierre J. Dufour, Sara Little, Douglas V. Guzior, John S. Petersen, James M. Shanahan, Bradley J. Carver

## Abstract

Obesity and diabetes are associated with worse prognosis for numerous malignancies. Both insulin resistance and altered levels of adipokines may explain the link between obesity and tumor progression. In preclinical models, METAP2 inhibitors induce weight loss and possess anti-tumor activity, but their effects on obesity-accelerated tumor growth are unknown. Here, we investigated the effects of SDX-7320, a novel polymer-conjugated METAP2 inhibitor, on obesity and obesity-accelerated tumor growth. The anti-obesity and metabolic effects of SDX-7320 were evaluated in diet-induced obese (DIO) mice. Pharmacokinetic-pharmacodynamic relationships for SDX-7320 and the active moiety SDX-7539, a fumagillin class METAP2 inhibitor, were assessed in DIO rats. Anti-tumor efficacy of SDX-7320 was assessed in syngeneic models of obesity-accelerated tumor growth. The anti-tumor efficacy of SDX-7320 and tirzepatide, a weight loss agent, were compared in DIO mice with MC38 tumors. Treatment with SDX-7320 stimulated weight loss in obese mice, increased insulin sensitivity, decreased plasma leptin, and increased plasma adiponectin. Pharmacokinetic-pharmacodynamic analysis showed greater anti-obesity efficacy in response to SDX-7320 than SDX-7539. SDX-7320 significantly attenuated obesity-accelerated tumor growth in three different models (B16F10, EO771, MC38). RNA-Seq analysis of MC38 tumors indicated that SDX-7320 suppressed expression of cell cycle genes (decreased G2M checkpoint and E2F target pathways) and increased expression of host immune response genes (elevated interferon alpha- and gamma-response pathways). In obese mice, SDX-7320 led to significantly greater MC38 tumor growth inhibition than tirzepatide, but caused less weight loss. Plasma metabolomics revealed non-overlapping effects of SDX-7320 and tirzepatide, consistent with different mechanisms of action. Taken together, we have shown for the first time that a METAP2 inhibitor attenuates obesity-accelerated tumor growth. Mechanistically, SDX-7320-mediated tumor growth inhibition likely results from both direct anti-tumor effects (given the observed intratumoral changes in the expression of cell cycle and immune response genes), and indirect effects on the host including weight loss, decreased adipose mass, improved insulin sensitivity and normalization of plasma leptin and adiponectin levels. The fact that SDX-7320 caused greater tumor growth inhibition than tirzepatide, yet caused less weight loss, suggests that direct anti-tumor effects significantly contribute to the anti-tumor activity of SDX-7320.

## Introduction

Obesity is associated with increased cancer incidence and worse prognosis for many tumor types (1). Both local and systemic factors, including genomic and transcriptomic alterations (2) and changes in the tumor microenvironment (3) have been proposed to explain the link between obesity and poor prognosis in cancer patients (4). Obesity causes subclinical inflammation in adipose tissues, contributing to the development of insulin resistance, which is linked, in turn, to cancer development and progression (5). In obese cancer patients, perturbations in circulating levels of insulin, adipose tissue-derived adipokines (e.g., leptin and adiponectin), and proinflammatory mediators (e.g., TNF-α, IL6) have been linked to tumor growth and metastasis (6, 7). Additionally, obesity is associated with increased expression of aromatase, the rate-limiting enzyme in estrogen biosynthesis, and reduced circulating levels of sex hormone-binding globulin, both of which have been suggested to contribute to poor outcomes in patients with estrogen-dependent tumors (8, 9). The abundance of mechanistic data linking obesity and systemic metabolic dysfunction to cancer progression has spawned the emergent field of *metabo-oncology*, which explores the biological link between deregulated host metabolism and tumor development and progression. In theory, novel therapeutics that target both tumors and systemic metabolic dysfunction should be clinically beneficial.

Methionine aminopeptidase type2 (METAP2) is a therapeutic target for various conditions including obesity and cancer (10). This metalloprotease plays a fundamental role in protein biosynthesis by catalyzing the removal of the N-terminal methionine residue from nascent polypeptides (10). A limited number of METAP2-specific substrates have been identified, including 14-3-3γ, cyclophilin A, eukaryotic elongation factors 1 and 2, thioredoxin-1, Rab-37, glyceraldehyde 3-phosphate dehydrogenase, hemoglobinβ and SH3 binding glutamic acid rich-like protein (11–13). Inhibition of the aminopeptidase function of METAP2 with fumagillin, a natural product secreted from the fungus *Aspergillus fumigatus Fresenius*, or fumagillin derivatives such as TNP-470 resulted in these substrates retaining their initiator methionine, affecting post-translational modifications, cellular localization, protein folding, and stabilization of these selective substrates (14). Fumagillin-derived METAP2 inhibitors stimulate weight loss in both non-clinical models of obesity and in obese patients (15–17). Weight loss mediated by METAP2 inhibition has been attributed to transient suppression of appetite, enhanced lipolysis in white adipose tissue, increased energy expenditure through direct action on brown adipocytes, and inhibition of angiogenesis in adipose tissue, with most of the weight loss due to reduction in adipose tissue as opposed to lean body mass (18–22). Mechanistically, the anti-cancer activity of METAP2 inhibitors has been attributed to inhibitory effects on cell proliferation and angiogenesis, among other pathways, such as MAPK and noncanonical Wnt signaling (15, 23–26).

While METAP2 is an attractive drug target for cancer therapeutics, the clinical utility of small molecule fumagillin derivatives has been limited by central nervous system (CNS) toxicity and poor drug-like properties (16, 27, 28). SDX-7320, a polymer-drug conjugate (PDC) of the novel METAP2 inhibitor SDX-7539, was developed to resolve these challenges by limiting CNS penetration, extending half-life, and increasing aqueous solubility, thereby improving overall safety while enabling less frequent administration (17). The efficacy of SDX-7320 against a variety of tumor types in lean mice and the results of a first-in-human dose-escalation safety and tolerability study (NCT02743637) have been previously described (17, 29).

This report describes a series of non-clinical studies investigating the ability of SDX-7320 to induce weight loss, alter host metabolic factors that have been implicated in obesity-related tumor progression and inhibit obesity-accelerated tumor growth. The main objectives of these studies were to 1) establish the effects of SDX-7320 on obesity and metabolic dysfunction in rodent models of diet-induced obesity (DIO); 2) determine whether SDX-7320 could attenuate obesity-accelerated tumor growth; and 3) define the potential mechanisms underlying the anti-tumor activity of SDX-7320.

## Materials and Methods

### Synthesis of SDX-7320

SDX-7320 was synthesized as previously described (29).

### *In Vivo* Studies

All animal experiments were carried out in accordance with the institutional guidelines and were approved by the Institutional Animal Care and Use Committee.

### Mouse DIO Model

Male C57BL/6j mice were individually housed and fed a high-fat, high-sucrose diet (Research Diets D12492) beginning at 5 weeks of age for 12 weeks prior to study initiation. Body weight was measured every four days, and the total food consumed per week per cage was also measured. Mice were randomized to treatment groups based on body weight (N=10/group) and treatment was conducted with subcutaneous (SC)-injected vehicle (5% w/v D-mannitol in water) Q4D or SDX-7320 (SC, Q4D) for a total of eight doses. Mice were euthanized by carbon dioxide inhalation on day 34 (two days after their last dose) and terminal blood was collected via cardiac puncture. Leptin and adiponectin levels were measured in plasma prepared from terminal blood samples using a Meso Scale Discovery mouse metabolic kit (K15124C) and adiponectin kit (K152BXC), respectively, according to the manufacturer’s instructions. Adipose tissue (epididymal, inguinal, and retroperitoneal depots) was dissected and weighed.

### Insulin Tolerance Test in DIO Mice

Male C57BL/6j mice (Jackson Laboratories) were individually housed and fed a high-fat, high-sucrose diet (Research Diets D12492) beginning at 6 weeks of age for 22 weeks before the start of the study. Body weight was measured on Day −4 and Day 0. Prior to treatment on Day −4, mice were randomized by body weight into vehicle and SDX-7320 treatment groups (N=6/group). The mice were then administered a single dose of vehicle (5% w/v D-mannitol in water) or SDX-7320 (8 mg/kg) SC. Four days later, on Day 0, baseline blood glucose (mg/dL) was measured (from <10 µL of blood collected from a tail nick) using an EasyMax V glucometer. Insulin tolerance test (ITT) was conducted after a 4-hour fast by delivering 0.75 U insulin/kg dosed intraperitoneally (IP). Blood glucose levels were measured at the indicated times following insulin administration using a hand-held glucometer (EasyMax V).

### Levin Rat Obesity Model

Male, diet-sensitive (DS) Sprague-Dawley/Levin rats (30) were obtained from Taconic at 3 weeks of age. Rats were individually housed and fed a high-fat/high-sucrose diet (Harlan TD.06414, 60% of calories from fat, 21% of calories from carbohydrate rodent diet) for 9 weeks until their average body weight was approximately 600 g, and they continued to receive this diet for the remainder of the study. Treatment groups (N=3/group) consisted of vehicle (phosphate-buffered saline, PBS), SDX-7320 (0.3, 1.0, 3.0 mg/kg; Q4D) and SDX-7539 (1.0, 3.0 mg/kg; Q4D) with all agents administered by SC injection for a total of 16 doses over 67 days. The body weight of each animal was measured every other day and food consumption was measured weekly. Blood samples for plasma SDX-7539 bioanalysis were collected on day 57, 1, 2, 4, and 24 h after dosing with SDX-7320 or SDX-7539. Adipose tissue was dissected (epidydimal, inguinal and retroperitoneal depots) and weighed at the end of the study (day 67, two days after the last dose of either test agent), following euthanasia by carbon dioxide inhalation.

### B16-F10 Syngeneic Melanoma Model

Male C57BL/6J mice (Jackson Laboratories) were fed either a high-fat diet (D124592) or a low-fat diet (D12450J) beginning at 6 weeks of age for 15 weeks prior to tumor cell inoculation and were maintained on the same diet for the entire study. The B16-F10 mouse melanoma tumor cell line was obtained from ATCC. Cultures were maintained in DMEM (Gibco, Invitrogen) supplemented with 5% fetal bovine serum and housed at 37°C in a humidified 5% CO_2_ atmosphere. The study was initiated by injecting 2 × 10^5^ B16F10 melanoma cells into the flank of lean and obese mice. When the tumors reached approximately 100 mm^3^, treatment with the test article was commenced (N=10/group). Animals were dosed with SDX-7320 or vehicle (5% D-mannitol in water) via SC intra-scapular injection every four days for a total of 5 doses. The study was terminated on day 17 and mice were euthanized using carbon dioxide inhalation, and adipose tissue was dissected and weighed.

### EO771 Syngeneic Triple Negative Breast Cancer (TNBC) Model

Female C57BL/6J mice that had undergone surgical ovariectomy at 4 weeks of age to mimic the postmenopausal state were obtained from the Jackson Laboratory at 6 weeks of age. The mice were then maintained for approximately 20 weeks on either a high-fat/high-sucrose diet (D12451) or a low-fat/high-sucrose diet (D12450B) until the start of the study. EO771 tumor cells (a model of TNBC, obtained from CH3 Biosystems) were implanted into the fourth mammary fat pad (1 × 10^5^/mouse; implantation was conducted after mice were anesthetized with 4% isoflurane and 1.2% oxygen) at approximately 26 weeks of age. Treatment with vehicle (5% w/v D-mannitol in water, SC, Q4D) or SDX-7320 (8 mg/kg, SC, Q4D) was initiated when the tumors were approximately 60 mm^3^ (N=9/group). All mice were euthanized on day 15 (after receiving a total of 4 doses) using carbon dioxide inhalation, after which the tumors and adipose tissues were excised.

### MC38 Syngeneic Colorectal Cancer Model

Male DIO and lean age-matched C57BL/6J mice (Jackson Laboratories) were fed a 60% kcal high-fat/high-sucrose diet (D12492) or a 10% kcal low-fat/high-sucrose diet (D12450B) beginning at 6 weeks of age for 13 weeks prior to study initiation. Mice were housed to 1-3 per cage for the duration of the study. Body weight was recorded twice weekly and changes in body weight were calculated relative to the baseline body weight at the start of the study. MC38 tumor cells (Kerafast, RRID:CVCLB288) were suspended in 50% PBS and 50% Matrigel before being implanted into the SC space of the right rear flank (2×10^5^/mouse). Tumor volume was measured twice weekly using wireless Mitutoyo UWAVE-T digital calipers in conjunction with UWAVE-R to digitally record the measurements. Tumor volume was calculated ((l × w^2^) × π/6) using Microsoft Office Excel software. Once the average tumor volume reached approximately 100 mm^3^, mice were randomized to treatment groups based on tumor volume. Treatment with vehicle (5% w/v D-mannitol in water, SC, Q4D) or SDX-7320 (6 mg/kg, SC, Q4D) was initiated for a total of five doses (N=10/group). Upon termination of the study (on day 17, one day following the last dose), the mice were euthanized by carbon dioxide inhalation, blood was collected via cardiac puncture, and serum and plasma were prepared. Tumor tissues were dissected, weighed, and placed in RNAlater tubes. The adipose tissues (epididymal, retroperitoneal, and inguinal) were dissected and weighed.

An additional study was conducted in which SC MC38 tumors (MC38 cells (Kerafast), 1 × 10^6^ cells/mouse) were established in DIO mice (Taconic, male C57Bl/6tac mice that had been fed a high-fat/high-sucrose diet (D12492) beginning at 6 weeks of age for 12 weeks). When the tumors reached approximately 60 mm^3^, treatment with vehicle, SDX-7320 (6 mg/kg, SC, Q4D), or tirzepatide (30 nmol/kg, SC, QD) was initiated (N=8/group). Body weight was measured every other day, and tumor volume was assessed twice per week. Upon termination of the study (eighteen days after initiation and two days after the last dose of SDX-7320 and one day after the last dose of tirzepatide; a total of 5 doses of SDX-7320 and 18 doses of tirzepatide were delivered), the mice were euthanized by carbon dioxide inhalation and adipose tissue depots were dissected and weighed.

### Statistical Analysis of In Vivo Data

Time-dependent data were analyzed using repeated measures two-way ANOVA (Geisser-Greenhouse correction with Sidak’s multiple comparisons test). For the analysis of endpoint data (i.e., adipose tissue mass, plasma biomarkers) non-parametric Kruskal-Wallis one-way ANOVA with Dunn’s multiple comparisons test was used. All data were analyzed as described using GraphPad v10.0.

### RNA isolation, sequencing and analysis

Total RNA from MC38 tumors was extracted using the Qiagen RNeasy Plus Universal Mini Kit following the manufacturer’s instructions (Qiagen). RNA samples were quantified using a Qubit 2.0 Fluorometer (Life Technologies), and RNA integrity was checked using a TapeStation (Agilent Technologies). The RNA sequencing library was prepared using the NEBNext Ultra RNA Library Prep Kit for Illumina (New England Biolabs), validated on TapeStation (Agilent Technologies), and quantified using a Qubit 2.0 Fluorometer (Invitrogen) and quantitative PCR (KAPA Biosystems). The sequencing library was clustered in one lane of a flow cell and loaded on an Illumina HiSeq instrument (4000 or equivalent), according to the manufacturer’s instructions. The sample was sequenced using a 2×150bp Paired End (PE) configuration at a depth of 3 x 10^7^ reads/sample. Image analysis and base calling were performed using HiSeq Control Software (HCS). Raw sequence data (.bcl files) generated were converted into fastq files and de-multiplexed using Illumina’s bcl2fastq 2.17 software. One mismatch was allowed for index sequence identification. Analysis was performed in the Watershed® Cloud Data Lab (CDL). Raw RNA sequence reads were aligned to the mouse reference genome (GRCm38/mm10) with Gencode vM10 annotations using STAR v2.7.5c. RNA-Seq data were uploaded to the GEO database (GSE295063). Principal component analysis (PCA) was performed using the PCA function in the Python package scikit-learn. Normalized gene expression was used as an input to identify the first 10 components of global gene expression. Differential gene expression analysis was performed using the R DESeq2 package version 1.40.2 (31). Gene set enrichment analyses (GSEA) were performed using the R clusterProfiler package, version 4.12.0 (32) and the mouse hallmark gene set collection from MsigDB v2024.1 (33). For this analysis, the Wald statistic calculated by DESeq2 analysis was used to rank genes for GSEA analysis.

### Procedure for SDX-7539 bioanalysis in support of rat pharmacokinetics

Rat plasma samples were subjected to protein precipitation followed by separation on a Waters Acquity UPLC system with a Waters Acquity BEH C18 reverse-phase column (50 mm × 2.1 mm) using an acetonitrile-water solvent gradient containing 0.1% formic acid. Detection was achieved using an AB Sciex API 5500 mass spectrometer with positive electrospray ionization in MRM mode, providing an acceptable concentration range for SDX-7539 of up to 500 ng/mL (lower limit of quantitation was 0.1 ng/ml).

### Metabolite extraction for untargeted metabolomics analysis

Murine plasma (50 µL) was transferred onto an Oasis solid-phase extraction (SPE) system (Waters). Quality control (QC) samples were generated by pooling 10 µL plasma from each sample. Next, 200 µL of 1:1 methanol:acetonitrile (MeOH:ACN) was added to each well and shaken for 1 min at room temperature at 360 rpm, followed by 10 min of incubation. Next, 150 µL 2:2:1 MeOH:ACN:water was added to each well and incubated again for 10 min. Polar metabolites were eluted into a 96-well collection plate using a positive pressure manifold. This procedure was repeated using 100 µL of 2:2:1 MeOH:ACN:H_2_O in the same collection plate. The polar eluates were then covered and stored at −80°C until Liquid Chromatography-Mass Spectrometry (LC/MS) analysis. The SPE plates from the polar metabolite extraction were then washed twice with 500 µL of 1:1 methyl *tert*-butylether:methanol to elute non-polar metabolites into a new collection plate using a positive pressure manifold. The combined eluates were dried under a stream of nitrogen at room temperature, reconstituted with 200 µL of 1:1 isopropanol:methanol, and stored at −80°C prior to LC/MS analysis.

### LC-MS/MS metabolite data collection

All untargeted metabolomics data were acquired using an Agilent Q-TOF 6546 mass spectrometer. The mobile phases for polar metabolite analysis were as follows: A) 20 mM ammonium bicarbonate, 0.1% ammonium hydroxide, 5% ACN, 2.5 mM medronic acid in water, and B) 95% ACN. Chromatography was performed using a HILICON iHILIC (P)-Classic (2.1 x 100 mm). A 4 µL injection volume was used for both positive and negative ionization modes. HILIC/MS was performed using the following linear gradient at a flow rate of 250 µL/min: 0-1 min: 90% B, 1-12 min: 90-35% B, 12-12.5 min: 35-25% B, 12.5-14.5 min: 25% B. The column was re-equilibrated with 20 column volumes of 90% B. Mass spectrometry analysis was performed with a mass range of 100-1700 Da and 1 scan/s. MS/MS data were acquired in a data-dependent iterative fashion with a 1.3 *m/z* isolation window.

The mobile phases for non-polar metabolite analysis were: A) 10 mM ammonium formate and 5 µM InfinityLab Deactivator Additive (Agilent) in 5:3:2 water:ACN:IPA, and B) 10 mM ammonium formate in 1:9:90 water:ACN:IPA. Reverse phase liquid chromatography (RPLC) was performed using a Waters Acquity HSS T3 (2.1 x 100 mm). A 2 µL injection volume was used when data were acquired in positive ion mode, and a 4 µL injection volume was used when data were acquired in negative ion mode. RPLC/MS was performed using the following linear gradient at a flow rate of 400 µL/min: 0-2.5 min: 15-50% B, 2.5-2.6 min: 50-57% B, 2.6-9 min: 57-70% B, 9-9.1 min: 70-93% B, 9.1-11 min: 93-96% B, 11-11.1 min: 96-100% B, 11.1-12 min: 100% B, 12-12.2 min: 100-15% B. The column was re-equilibrated for 3.8 min. Mass spectrometry analysis was performed with a mass range of 100-1700 *m/z* and 1 scan/s. MS/MS data were acquired in a data-dependent iterative fashion with a 1.3 *m/z* isolation window.

### Untargeted metabolomics data processing and statistical analysis

Data files were converted to the mzML format using MSConvert. Features were detected using proprietary peak detection and curation software (Panome Bio, Inc). Features were aligned across samples, and those with intensities greater than 1/3 of the corresponding intensity in the QC sample were classified as contaminants and were removed from future analysis. Feature degeneracies (isotopes, adducts, fragments, etc.) were identified using clustering and ion assignment. Metabolites were structurally identified by comparing isotope patterns and MS/MS fragmentation data (when available) against a database composed of known metabolites found in RefMet, LipidMaps, and HMDB, in addition to an in-house standard library from Panome Bio (34). Metabolite signals were discarded if the coefficient of variation among the QC samples was greater than 25%. Missing values were imputed using half the minimum intensity detected for each metabolite.

Principal component analysis was performed using the PCA function within the Python package scikit-learn. Prior to hypothesis testing, data were log-transformed to ensure normality. Metabolites showing significant differences between the control, tirzepatide, and SDX-7320 treated mice were identified using one-way ANOVA.

#### Data availability statement

The data for this study were generated by multiple contract research organizations. Reports containing raw and derived data supporting the findings of this study are available from the corresponding author upon reasonable request. Gene expression data are available in the GEO database under accession number GSE295063.

## Results

### SDX-7320 stimulates weight loss and improves insulin sensitivity in obese mice

Small molecule METAP2 inhibitors exhibit anti-obesity and anti-diabetes activities, both in non-clinical models and in humans (20, 35–37). Therefore, the activity of SDX-7320, the polymer-conjugated METAP2 inhibitor, was evaluated in standard rodent models of DIO and insulin resistance. Treatment with SDX-7320 SC led to a dose-dependent reduction in body weight in DIO mice (Fig. 1A). Consistent with the initial weight loss mediated by SDX-7320, food intake (normalized to body weight) was significantly reduced during the first week of the study, but then increased, returning to vehicle group levels after the second week despite continued weight loss (Fig. 1B). Analysis of adipose tissue weight at the end of the study demonstrated a significant reduction in response to SDX-7320 in each of the three adipose depots (Fig. 1C-E). Mice treated with SDX-7320 had significantly reduced leptin levels (Fig. 1F) and increased adiponectin levels (Fig. 1G), which resulted in a significant reduction in the leptin-to-adiponectin ratio (Fig. 1H).

**Figure 1.**
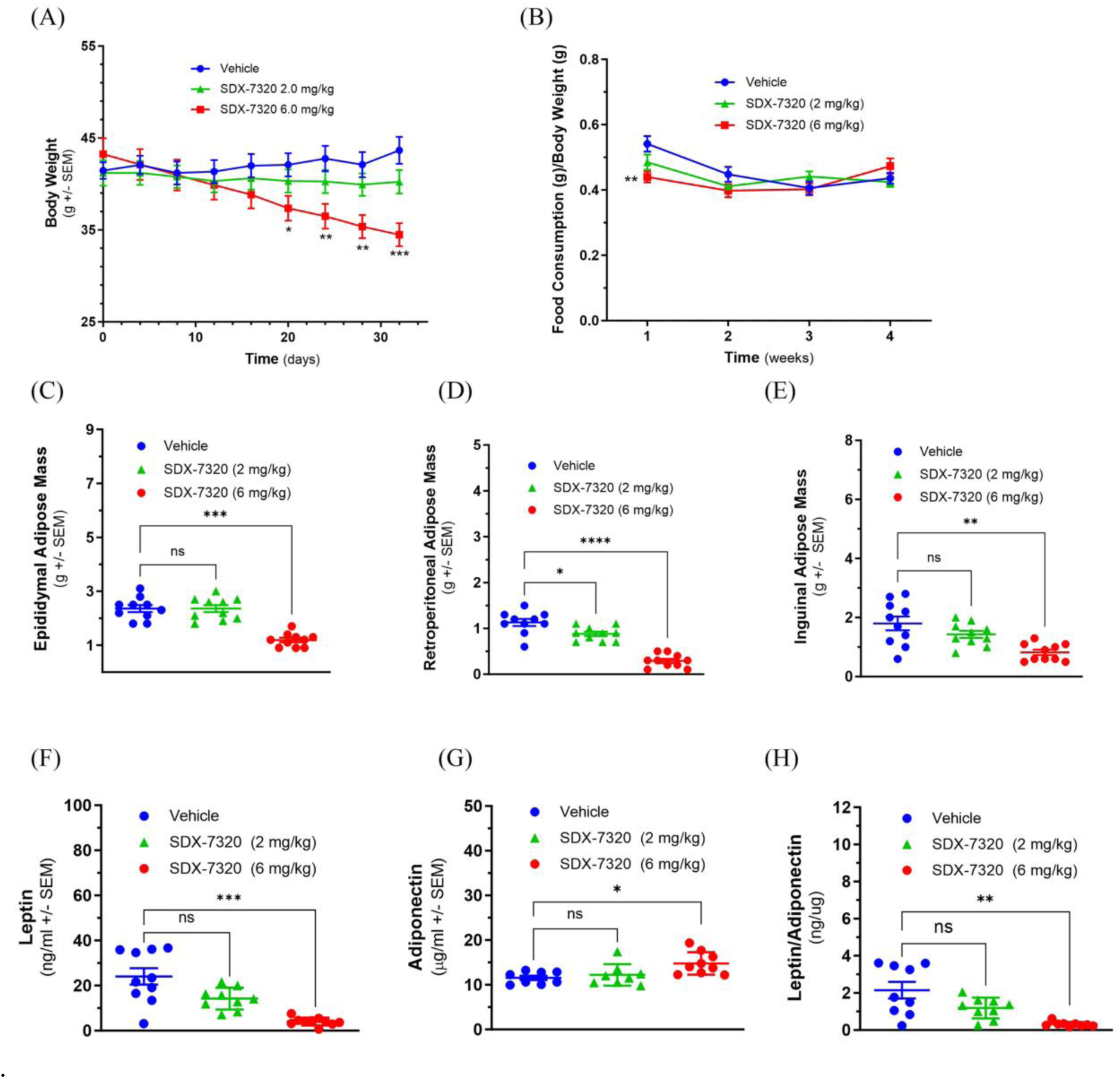
Effect of SDX-7320 on body weight, fat mass and adipokine levels in diet-induced obese mice. Diet-induced obese mice (male C57Bl/6j, n=6 group) were dosed subcutaneously with vehicle (5% mannitol in water) or with the indicated dose levels of SDX-7320 (Q4D dissolved in vehicle). Body weight was measured every four days (Panel A), and food consumption was measured weekly (Panel B). Thirty-three days after initiating dosing, mice were euthanized, adipose tissues were dissected and weighed (Panel C-E), and blood was collected for analysis of leptin and adiponectin (Panels F-H). Statistical analysis was carried out as described in the methods section (*p < 0.05, **p < 0.01, ***p < 0.005, ****p <0.001 versus vehicle-treated mice).

In a second study in DIO mice, SDX-7320 elicited effects on body weight, food intake, and fat mass similar to those shown in Figure 1 (Supplementary Fig. S1). To assess the impact of weight loss in response to SDX-7320 on glycemic control, an intraperitoneal glucose tolerance test (IPGTT) was conducted after four weeks of SDX-7320 treatment. IPGTT results showed a significant dose-dependent reduction in glucose excursion, indicating enhanced glucose disposal in mice treated with SDX-7320 (Supplementary Figs. S1D, E). Consistent with the reductions in body weight and fat mass in response to SDX-7320, decreases in plasma levels of ALT, cholesterol, triglycerides, as well as in liver weight, were observed (Supplementary Figs. S1F-I).

Because DIO mice treated with SDX-7320 exhibited improved glucose responses in an IPGTT, a separate study was conducted to directly assess whether SDX-7320 improved peripheral insulin sensitivity. An ITT was conducted in DIO mice that received vehicle or a single dose of SDX-7320 four days before the ITT. The acute response to insulin (i.e., a decline in blood glucose) in mice pretreated with SDX-7320 was significantly greater than that observed in mice pretreated with the vehicle (Supplementary Fig. S2). These results demonstrated that SDX-7320 enhances peripheral insulin sensitivity.

### Effect of polymer conjugation on anti-obesity activity of the METAP2 inhibitor SDX-7539

SDX-7320 was designed to minimize CNS toxicity previously observed with small-molecule METAP2 inhibitors, such as TNP-470, by attaching a novel small-molecule METAP2 inhibitor (SDX-7539) to a high-molecular-weight, water-soluble, and biocompatible polymer. Pharmacokinetic studies of unconjugated SDX-7539 versus SDX-7539 derived from the PDC SDX-7320 showed that the half-life of SDX-7539 in plasma was approximately 40-fold longer following administration of the PDC (29). To investigate whether the unique plasma PK profile observed for SDX-7539 delivered via the polymer conjugate had any effect on its pharmacodynamics, we compared the anti-obesity activity of SDX-7320 with that of unconjugated SDX-7539 in obese male Levin rats. Subcutaneous administration of SDX-7320 elicited a dose-dependent attenuation in weight gain over time (Fig. 2A), accompanied by a reduction in adipose tissue (Supplementary Fig. S3), and improved insulin sensitivity (Supplementary Fig. S3). Significant differences in SDX-7539 C_max_ and AUC were observed; dosing with SDX-7320 yielded far lower systemic plasma levels of SDX-7539 than dosing with unconjugated SDX-7539 (Fig. 2B, Supplementary Table S1). PK-PD analysis of SDX-7539 pharmacokinetics (C_max_, AUC) against the efficacy data (i.e., % increase from baseline in body weight) showed greater anti-obesity efficacy in response to SDX-7320, despite much lower plasma exposure to polymer-released SDX-7539 (Figs. 2C, D).

**Figure 2.**
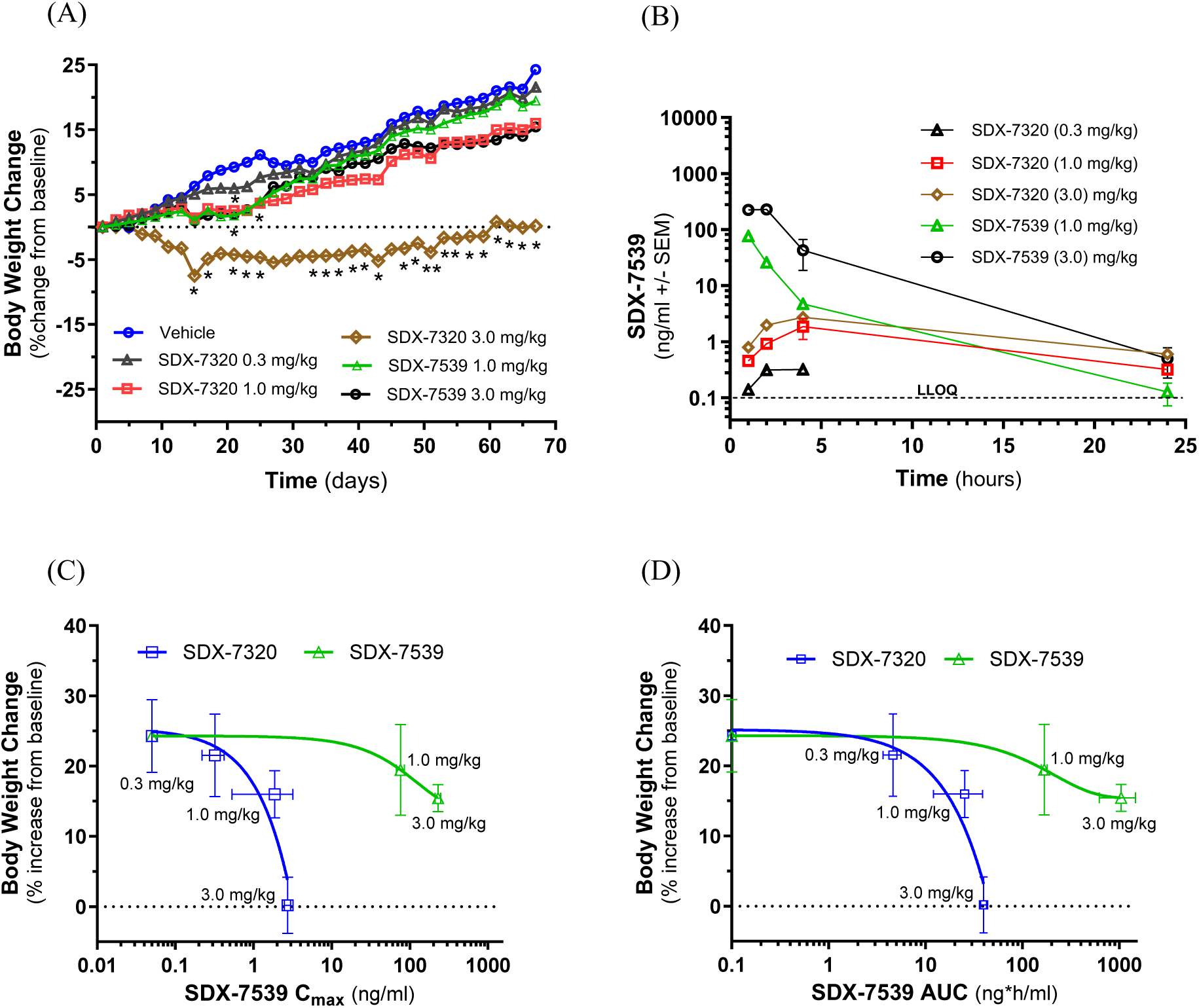
PK-PD analysis of SDX-7320 compared to SDX-7539 in obese rats. Obese, male Sprague-Dawley (Levin) rats were dosed (via subcutaneous injection; Q4D; N=3/group) with vehicle (PBS), SDX-7320 (0.3, 1.0, 3.0 mg/kg) or SDX-7539 (1.0, 3.0 mg/kg). Body weight was measured every other day and normalized to baseline values (Panel A). On day 56 of the study, at 1-, 2-, 4-, and 24-hours post-dose, blood samples were collected, plasma was prepared and analyzed by LC/MS for levels of SDX-7539 (Panel B). Pharmacokinetic parameters (C_max_, AUC) were calculated from the SDX-7539 exposure data (see Supplementary Table S1) and plotted versus the % increase in body weight from baseline (Panels C, D). Data were analyzed as described in the methods section (*p < 0.05, **p < 0.01 versus vehicle-treated rats). The PK-PD data presented in panels C and D were analyzed by first calculating the AUC for each group with significance assessed using unpaired t-tests (the results in panels C and D comparing vehicle-treated and SDX-7320-treated animals were significantly different, with *p < 0.05).

### SDX-7320 inhibits obesity-accelerated tumor growth

Having established the anti-obesity activity of SDX-7320 including its effects on host metabolic factors (insulin sensitivity, leptin and adiponectin levels), we next determined its ability to suppress obesity-accelerated tumor growth. Three different mouse tumor models were selected because obesity is known to accelerate the growth of each of these tumors (38–40). Initially, the B16F10 melanoma model was used to evaluate the effects of SDX-7320 on obesity-accelerated tumor growth. Subcutaneous B16F10 tumors grew significantly faster and achieved a larger size in obese animals than in lean age-matched animals (Fig. 3A). SDX-7320 significantly inhibited tumor growth in both obese and lean animals in a dose-dependent manner (Figs. 3B, D). SDX-7320 significantly reduced absolute body weight in both obese and lean mice compared to their respective vehicle-treated controls (Figs. 3C, E). In obese mice, weight loss induced by SDX-7320 resulted in part from significant reduction in adipose tissue mass (Fig. 3F-H). SDX-7320 did not significantly reduce adipose tissue mass in lean mice bearing B16F10 tumors (Fig. 3F-H).

**Figure 3.**
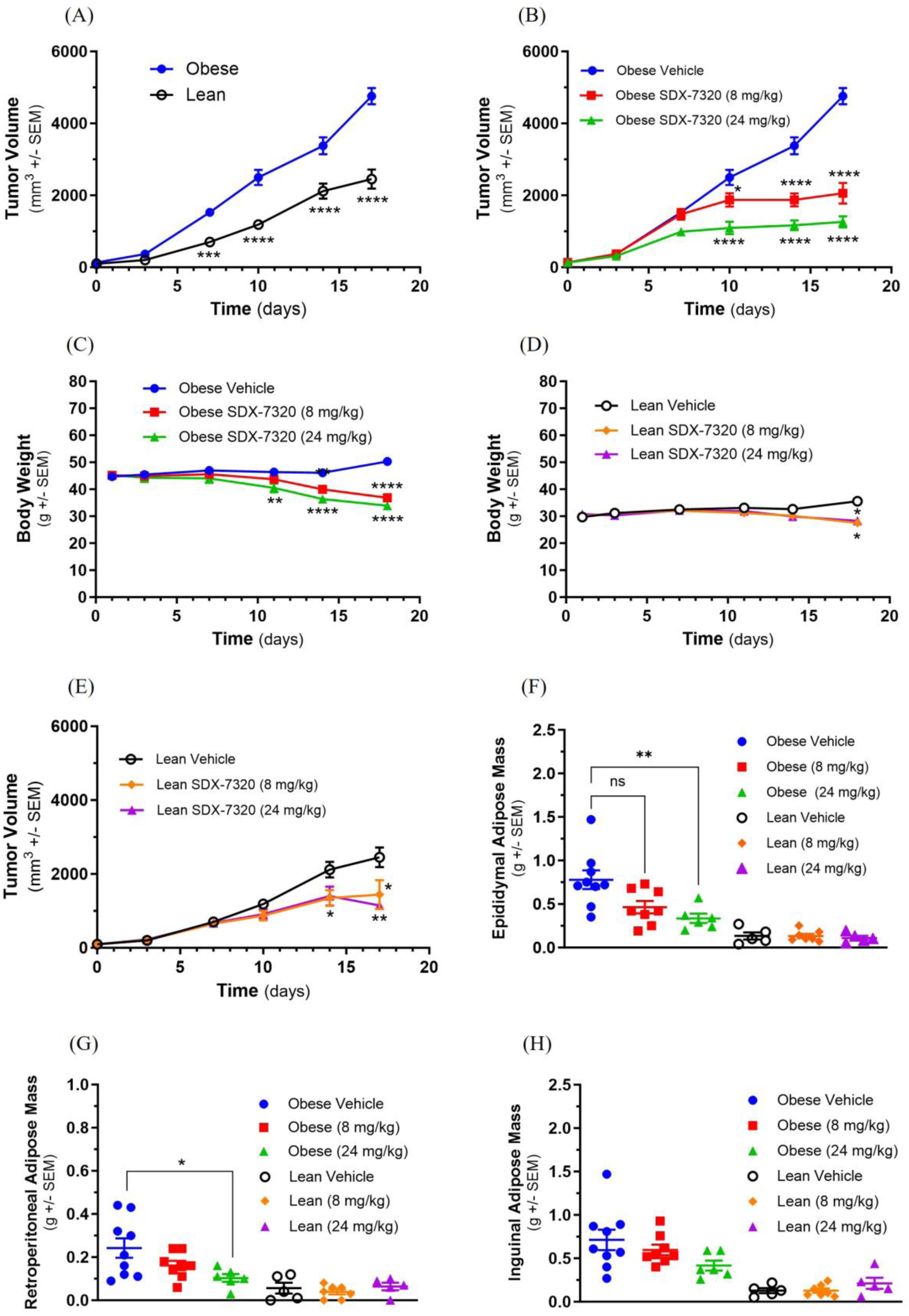
SDX-7320 inhibits the growth of obesity-accelerated B16F10 melanoma. Diet-induced obese mice (male C57Bl/6j, n=10 group) or their lean littermates (n=10/group) were injected with B16F10 melanoma cells into the subcutaneous space of the right rear flank (1×10^5^ cells/mouse; 50/50 Matrigel/PBS). When tumors were approximately 100 mm^3^, mice were dosed subcutaneously with vehicle (5% mannitol in water) or with the indicated dose levels of SDX-7320 (Q4D dissolved in vehicle). Tumor volume was measured every fourth day (Panels A, B, and D), body weight was measured twice/week (panels C, E). Seventeen days after initiating dosing, mice were euthanized and adipose tissues dissected and weighed (panels F-H). Data were analyzed using two-way ANOVA with multiple comparisons for results in panels A-E and one-way ANOVA for results in panel F-H as described in the methods section (*p < 0.05, **p < 0.01, ***p < 0.005, ****p <0.001 versus vehicle-treated mice).

To further investigate the efficacy of SDX-7320 in obesity-accelerated tumor growth, a study was conducted in obese vs. lean ovariectomized female C57BL/6 mice (intended to mimic the post-menopausal state) harboring syngeneic, orthotopic EO771 mammary gland tumors. The mice were fed either a high-fat diet (DIO mice) or a low-fat diet (lean mice). As shown in Supplementary Fig. S4A, tumor growth was significantly accelerated in obese compared to lean mice. Obese and lean mice bearing EO771 mammary gland tumors were randomized to treatment with vehicle or SDX-7320 (8 mg/kg, SC, Q4D) once the mean tumor volume had reached approximately 60 mm^3^. Treatment with SDX-7320 significantly reduced the rate of tumor growth in obese mice (Supplementary Fig. S4B), whereas only a trend was observed in lean mice (Supplementary Fig. S4D). There was also a trend towards reduced body weight with SDX-7320 in obese mice with EO771 tumors, and no effect on body weight was observed in lean mice with EO771 tumors (Supplementary Figs. S4C, E). SDX-7320 significantly reduced the mass of retroperitoneal, inguinal and parametrial adipose tissue in obese mice with trends to reduction of retroperitoneal and parametrial adipose tissue in lean mice (Supplementary Fig. S4F-H).

To extend the findings described above in models of obesity-accelerated melanoma and breast cancer, the effect of SDX-7320 on MC38 tumor growth, a model of colorectal cancer, was tested. Male C57BL/6 mice were initially fed either a high-fat/high-sucrose diet or a low-fat diet for 12 weeks to generate obese and lean mice, respectively. Subsequently, obese and lean mice were injected SC with MC38 tumor cells and tumor growth was monitored. Consistent with the models mentioned above, tumor growth in obese mice was significantly greater than that in lean mice (Fig. 4A). Treatment with SDX-7320 was initiated when the average tumor volume reached approximately 100 mm^3^.

**Figure 4.**
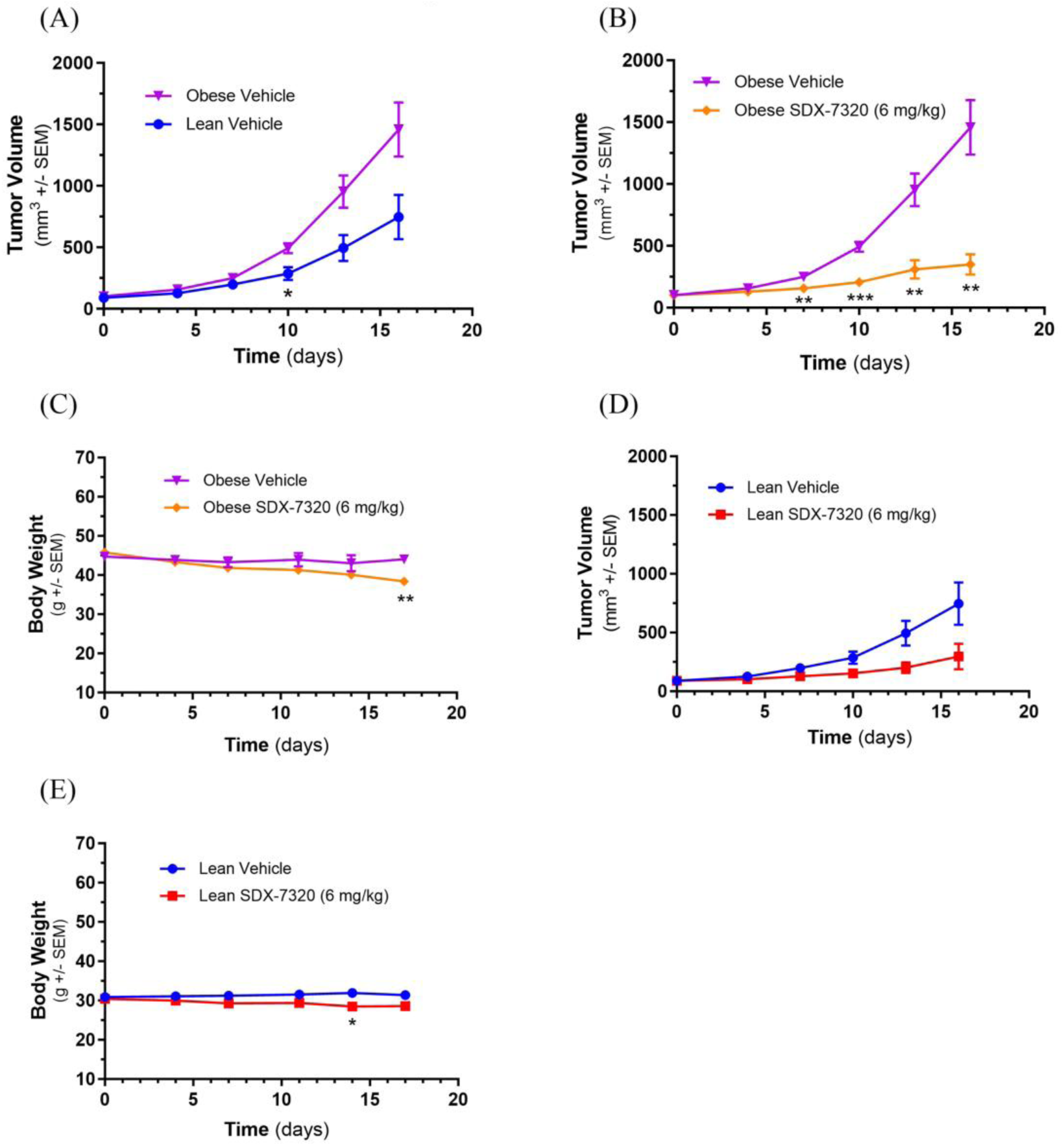
SDX-7320 inhibits the growth of obesity-accelerated MC38 tumors. Diet-induced obese mice (male C57Bl/6, n=10/group) or their lean littermates (n=10/group) were injected in the right rear flank with MC38 tumor cells (2 x 10^5^ cells/mouse). When tumors were approximately 100 mm^3^, mice were dosed subcutaneously with vehicle (5% mannitol in water) or with SDX-7320. Body weight (Panels C and E) and tumor volume (Panels A, B and D) were measured twice/week. In-life data were analyzed using two-way ANOVA with multiple comparisons as described in the methods section (*p < 0.05, **p < 0.01, ***p <0.005 versus vehicle-treated mice).

SDX-7320 treatment significantly inhibited tumor growth in obese mice whereas a trend towards reduction in tumor growth was observed in lean mice (Figs 4B, D). SDX-7320 elicited modest but significant reduction in body weight in cohorts of obese and lean mice (Figs. 4C, E).

To investigate whether there were distinct alterations in gene expression across the different cohorts and to elucidate the mechanisms underlying SDX-7320-mediated inhibition of MC38 tumor growth in both obese and lean mice, RNA sequencing was conducted on tumor tissue. A total of 463 up-regulated and 61 down-regulated differentially expressed genes (DEGs; p.adj<0.05, fold change>1.5) were found in tumors from obese mice treated with SDX-7320 vs. vehicle (Fig. 5A, B; Supplementary Table S2; Supplementary Fig. S5) using DESeq2. In tumors from lean mice, SDX-7320 treatment was associated with only 14 up-regulated and 11 down-regulated DEGs (Figs. 5A, B; Supplementary Table S2; Supplementary Fig. S5). To better understand the significance of these global transcriptome differences, pathway enrichment analysis was performed. The results of ranked gene set enrichment analysis (GSEA) showed that the expression of numerous pathways was enriched in tumors from SDX-7320 vs. vehicle-treated obese mice (Fig. 5C), including the immune response elements interferon gamma response, interferon alpha response, inflammatory response, IL6-JAK-Stat3 signaling, and TNF-α signaling via NFKappaB. The negatively enriched pathways included G2M checkpoint and E2F target pathways. In lean mice, fewer pathways were positively enriched by SDX-7320 treatment, although two of the same immune response elements were enriched (i.e., interferon alpha response and interferon gamma response; Fig. 5D), but not IL6-JAK-Stat3 or TNF-α signaling. Negatively enriched pathways in tumors from lean mice treated with SDX-7320 also included G2M checkpoint and E2F target pathways, as was observed in obese mice. The responses in lean mice generally showed lower enrichment scores and less robust false discovery rates than those observed in obese mice. As multiple immune response pathways were enriched in SDX-7320-treated tumors from both obese and lean mice, we next determined whether there were any overlapping effects of SDX-7320 on specific DEGs. Venn diagrams for SDX-7320-treated groups vs. vehicle treatment indicated that there were 10 shared up-regulated and 3 shared down-regulated DEGs in tumors from obese and lean mice treated groups (p.adj<0.05, fold change >1.5; Figs. 5A, B). The overlapping up- and down-regulated genes suggest a consistent effect of METAP2 inhibition on tumor gene expression regardless of body weight (highlighted in orange and purple, respectively, in Supplementary Table S2). For example, the expression of numerous granzymes and perforin1 was increased in tumors from both obese and lean mice treated with SDX-7320 (Fig. 5E-5G), suggesting that inhibiting METAP2 enhanced the host anti-tumor immune response. Furthermore, the expression of CD-206/MRC1, a marker of immune-suppressive macrophages, was significantly decreased in tumors from both lean and obese mice, suggesting that SDX-7320 favorably alters the tumor immune microenvironment (Fig. 5H).

**Figure 5.**
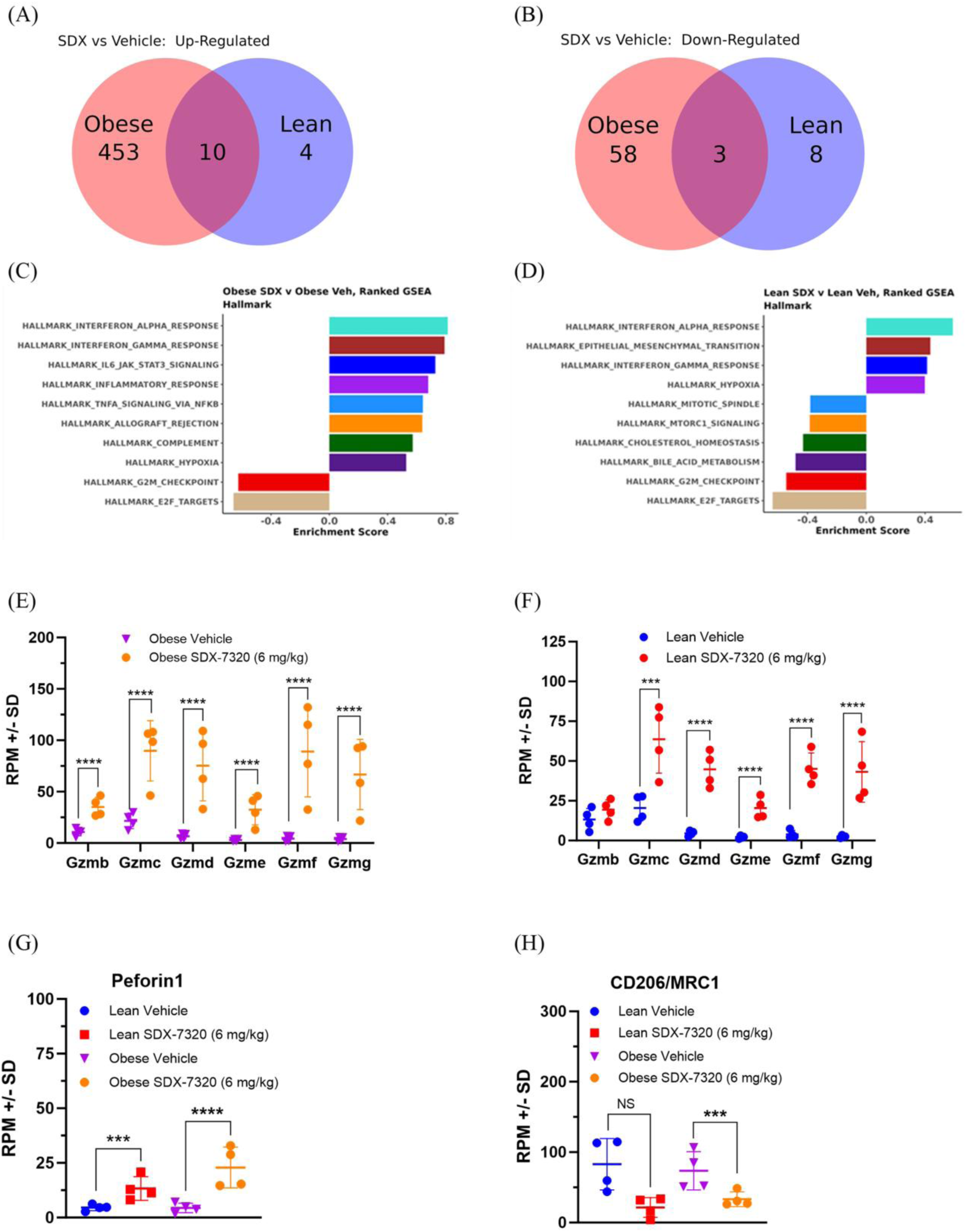
Effect of SDX-7320 on gene expression in MC38 tumors from obese vs. lean mice. Diet-induced obese or lean male mice were injected in the right rear flank with MC38 tumor cells. When tumors were approximately 100 mm^3^, mice were dosed subcutaneously with vehicle (5% mannitol in water) or with SDX-7320 (6 mg/kg). Upon study termination, tumor tissue was placed into RNALater (n=4/group), after which bulk RNA-Seq was carried out. Venn diagrams showing DEGs for SDX-7320-mediated up-regulated (Panel A) and down-regulated (Panel B) genes in MC38 tumors from obese and lean mice. Gene set enrichment analyses for SDX-7320 vs. vehicle-treatments are shown for tumors harvested from obese (Panel C) and lean (Panel D) mice, respectively. The effect of SDX-7320 vs. vehicle on selected differentially expressed genes including granzyme mRNA levels (Panels E and F), perforin1 mRNA levels (Panel G) and CD-206/MRC1 mRNA levels (Panel H) from obese and lean mice are shown (adjusted p values ***p < 0.005, ****p <0.001 versus vehicle-treated mice).

### Effects of SDX-7320 versus tirzepatide on MC38 tumor growth in DIO mice

With an increasing focus on the link between obesity and cancer progression and the clinical success of incretin-based therapies for weight loss and improvement of obesity-related conditions, we compared the anti-tumor efficacy of SDX-7320 with tirzepatide in the MC38 syngeneic model of obesity-accelerated colon cancer. DIO mice with established MC38 tumors were treated with vehicle, SDX-7320 or tirzepatide beginning when the tumors were approximately 60 mm^3^. Interestingly, tirzepatide modestly but significantly inhibited MC38 tumor growth, whereas SDX-7320 exerted a significantly greater inhibitory effect on MC38 tumor growth compared to tirzepatide (Fig 6A). In contrast, the effect of tirzepatide on body weight loss was significantly greater than that of SDX-7320 (Fig. 6B). Unsurprisingly, when adipose tissue was measured at the end of the study, the effect of tirzepatide on fat mass was greater than that of SDX-7320 (Fig. 6C-E). Analysis of circulating adipokines leptin and adiponectin showed that tirzepatide elicited a greater reduction in plasma leptin levels than SDX-7320 (Fig. 6F), whereas only SDX-7320 significantly increased levels of plasma adiponectin (Fig. 6G). The leptin:adiponectin ratio (LAR) was determined and only tirzepatide significantly decreased LAR (Fig. 6H).

**Figure 6.**
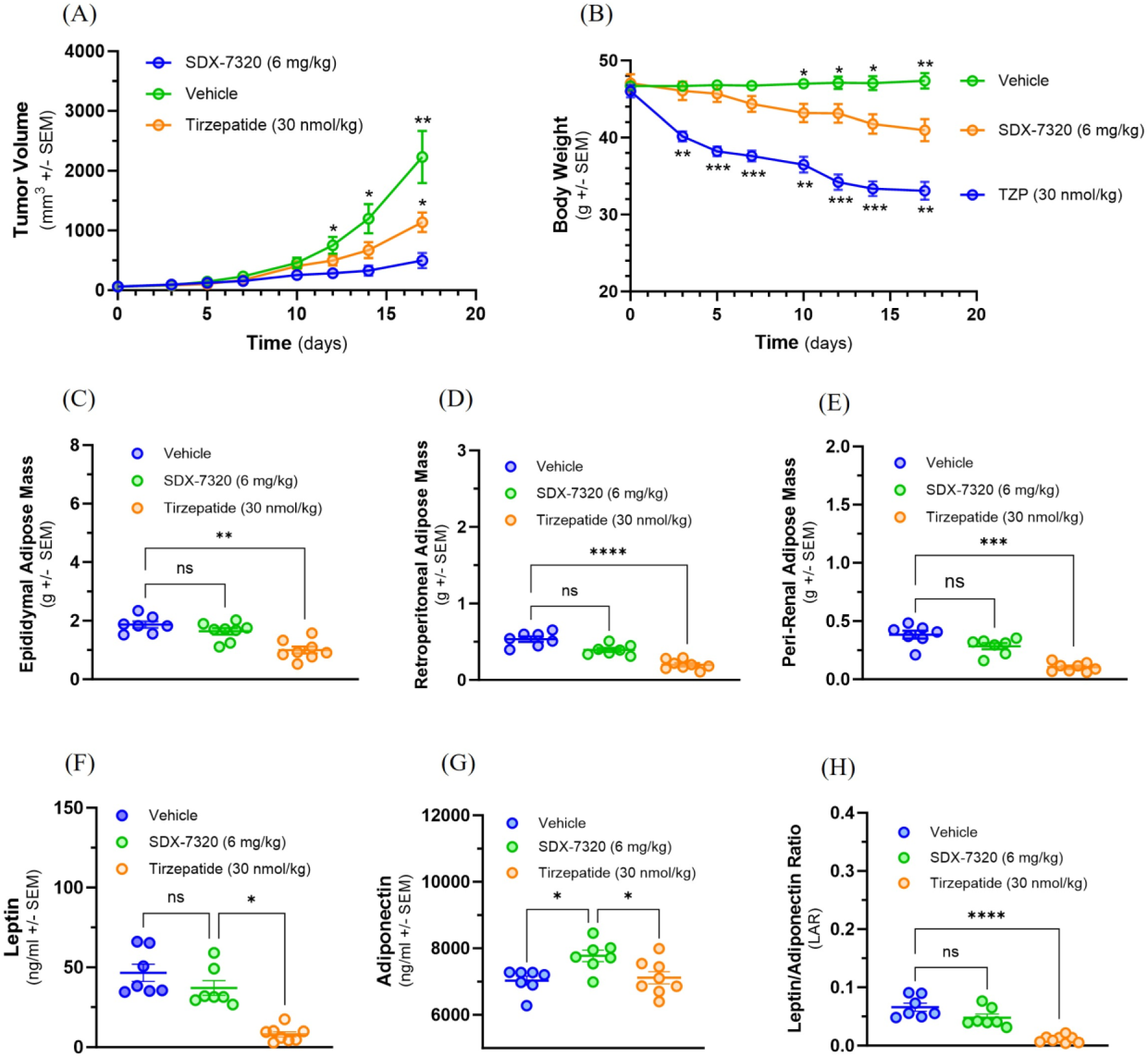
Comparison of anti-tumor activity of SDX-7320 and tirzepatide on MC38 tumor growth in obese mice. Diet-induced obese mice (male C57Bl/6j, n=8/group) were injected in the right rear flank with MC38 tumor cells (1 x 10^6^ cells/mouse). When tumors were approximately 60 mm^3^, mice were dosed subcutaneously with vehicle (5% mannitol in water), SDX-7320 (6 mg/kg, Q4D) or tirzepatide (30 nmol/kg, QD). Tumor volume and body weight were measured twice/week (Panels A and B). Eighteen days after initiating dosing, animals were euthanized, and adipose tissue from different depots was dissected and weighed (Panel C-E). Plasma was prepared for analysis of adipokines leptin and adiponectin (Panels F-H). Data were analyzed as described in the methods section (*p < 0.05, **p < 0.01, ***p < 0.005, ****p <0.001). Global untargeted metabolomics data were analyzed using principal component analysis (Panel H). Metabolites (n=5) from SDX-7320 and tirzepatide treatment groups were significantly different from the vehicle-treated group (Panel I).

To explore potential mechanistic similarities and differences between SDX-7320 and tirzepatide, global untargeted metabolomics was performed on plasma samples collected at the end of the study. The overall chemical profiles of each sample were investigated using PCA, which revealed a clear separation between the treatment groups (Fig. 6H), suggesting non-overlapping mechanisms of action. Treatment with tirzepatide resulted in increased levels of six metabolites and decreased levels of 22 metabolites compared to vehicle controls (Supplementary Table S3). Treatment with SDX-7320 resulted in 15 metabolites showing significant alterations, of which four were significantly up-regulated and 11 were significantly down-regulated, whereas only five of the metabolites were significantly altered by both treatments (Supplementary Table S3; Supplementary Fig. S6). One steroid, ST 24:1;O5, was up-regulated by both compounds, whereas dihydrouracil, ureidopropionic acid, and diacylglycerophosphocholines PC 44:10 and PC 40:6 were down-regulated by both agents (Fig. 6I). Changes in the remaining 10 compounds exclusively altered by SDX-7320 included increases in artemin and glycerophosphoglycerols PG O-36:5 and PG 34:1, in tandem with the depletion of several additional lipids (Supplementary Table S3). Interestingly, the depleted lipid profile included ceramides and an additional phosphatidylcholine (PC O-38:4). Lastly, investigating the significantly altered metabolites exclusive to tirzepatide showed that most down-regulated metabolites were phospholipids and di/triglycerides (Supplementary Table S3).

## Discussion

In 2003, Calle et al. reported that obesity was associated with worse clinical outcomes in more than 10 tumor types. Since then, numerous studies have reported that obesity and systemic metabolic dysfunction (e.g., insulin resistance, metabolic syndrome, and type 2 diabetes) play important, yet incompletely understood roles in cancer progression, response to therapy, and patient survival (41–43). Here we have shown in obese mice that treatment with the novel PDC METAP2 inhibitor SDX-7320 led to weight loss, improved metabolic function and significant inhibition of tumor growth. To our knowledge, this is the first demonstration that a METAP2 inhibitor reduces obesity-accelerated tumor growth.

SDX-7320 reduced adipose mass, increased insulin sensitivity and altered adipokine levels in rodent models of obesity and metabolic dysfunction. In obese rats, the polymer conjugate SDX-7320 was superior in attenuating body weight gain compared to the unconjugated small molecule METAP2 inhibitor SDX-7539. Polymer conjugation of the METAP2 inhibitor SDX-7539 conferred a pharmacokinetic and pharmacodynamic advantage over the unconjugated inhibitor, suggesting an improved therapeutic index of SDX-7320 compared to small-molecule METAP2 inhibitors. The reduction in body weight and in adipose tissue mass in obese animals after treatment with SDX-7320 was partly due to a transient reduction in food consumption. Interestingly, food intake returned to normal during the course of the study despite sustained body weight reduction while on SDX-7320, suggesting that additional mechanisms beyond inhibition of caloric intake contribute to the maintenance of reduced body weight. Notably, as mentioned above, the activation of brown adipose tissue (and the resultant increase in metabolic rate) and inhibition of angiogenesis in adipose tissue have previously been reported to contribute to the anti-obesity effect of METAP2 inhibitors (19, 21). Interestingly a methionine-restricted diet has shown effects on metabolic endpoints in obese mice quite similar to what we have observed in response to SDX-7320, raising the possibility that some of the effects downstream of METAP2 inhibition may result from decreases in intracellular methionine (44). Regardless of the mechanism by which SDX-7320 induces weight loss and improves insulin sensitivity in obese mice, its effect on plasma leptin (decreased) and adiponectin (increased) may also have contributed to tumor growth inhibition, given the evidence that these adipokines exert opposing effects on tumor growth (45, 46). In addition to indirect endocrine effects on tumor growth, SDX-7320 also had direct anti-tumor effects. In both obese and lean mice with syngeneic MC38 colorectal tumors, RNA-seq data indicated that SDX-7320 negatively enriched the G2M checkpoint and E2F transcription factor target pathways, which is consistent with prior evidence that METAP2 inhibitors suppress the proliferation of tumor cells via cell cycle arrest (47, 48). These RNA-seq findings provide new insights into the mechanism underlying the ability of METAP2 inhibitors to suppress the proliferation of tumor cells. Interestingly, the key hallmarks of an anti-tumor immune response were observed, including a significant increase in the expression of genes encoding granzymes and perforin1, suggesting that treatment with SDX-7320 may have enhanced the infiltration of immune cells and their cytotoxic activity within the tumors. Furthermore, decreased expression of MRC-1/CD206 was observed in tumors from mice treated with SDX-7320, suggesting reversal of an immunosuppressive tumor microenvironment.

There is significant interest in targeting excessive weight and metabolic dysfunction as a complementary treatment for cancer. Initial clinical exploration of intentional weight loss in patients with cancer has shown modest success, as demonstrated in studies by the Women’s Health Initiative (49, 50). Dietary interventional clinical trials (e.g., calorie restriction and a fasting-mimicking diet) in combination with chemotherapy have shown improved clinical outcomes for patients with a variety of cancer types (51), although real-world compliance with diets is challenging in this patient population. The anti-diabetic drug metformin has been investigated as a pharmacological intervention to treat cancer patients based on findings in observational studies and its ability to inhibit tumor growth in non-clinical experiments (52). Recently, the MA.32 clinical trial tested the effect of metformin on cancer outcomes in a broad selection of women with hormone receptor-positive breast cancer. However, the primary endpoint of invasive disease-free survival was not met, possibly because of metabolic heterogeneity resulting from the unselected patient study design (53).

New obesity drugs based on incretin hormones such as glucagon-like peptide 1 (GLP-1) and gastric inhibitory polypeptide (GIP) have transformed the landscape of obesity therapeutics. Anti-obesity drugs are now being evaluated as possible preventative and/or therapeutic agents in non-clinical models of obesity-accelerated cancer as well as in cancer patients (54, 55). Given the anti-tumor, anti-obesity and insulin-sensitizing attributes of SDX-7320, we compared one of these anti-obesity drugs (tirzepatide) to SDX-7320 in obese mice bearing MC38 tumors. In comparison to tirzepatide, treatment of obese tumor-bearing mice with SDX-7320 led to significantly greater anti-tumor effects but less weight loss. These results raise the possibility that the direct anti-tumor effects of METAP2 inhibition contribute more than its indirect effects on host metabolism (e.g., weight loss, insulin sensitization) to explain superior tumor growth inhibition relative to tirzepatide. However, one could anticipate that the relative importance of direct vs. indirect (host) anti-tumor effects of SDX-7320 may vary in different tumor types in humans versus mice. To more fully understand the overall anti-tumor effects of SDX-7320, additional experiments are warranted to determine the relative importance of direct anti-tumor effects compared to indirect metabolic effects.

Because treatment with both SDX-7320 and tirzepatide led to weight loss and inhibition of tumor growth (albeit to differing degrees), it was important to evaluate the effects of these agents on plasma metabolites. Consistent with having different mechanisms of action, treatment with SDX-7320 or tirzepatide led to a series of unique and non-overlapping changes in metabolite levels. For example, treatment with SDX-7320 led to significantly lower plasma ceramide levels including hexosyl-ceramide whereas treatment with tirzepatide did not have this effect (Supplementary Table S3). Hexosyl-ceramide has been associated with cancer cell drug resistance and metastasis (56, 57) whereas ceramides have been widely studied as potential mediators of inflammation and insulin resistance (58). Treatment with SDX-7320 or tirzepatide led to overlapping changes in the levels of several plasma metabolites (Supplementary Table S3). For example, both treatments led to decreased levels of dihydrouracil and ureiodopropionic acid, which are products of uracil catabolism. Ureiodopropionate is the breakdown product of dihydrouracil and is produced by the enzyme dihydropyrimidase (59). Whether altered uracil metabolism helps to explain either the anti-tumor or weight loss effects of these two agents is uncertain and warrants further investigation. Based on these preclinical findings in obese tumor bearing mice, it will be of significant interest to determine whether similar effects on metabolites occur in obese cancer patients.

Overall, this study has a number of strengths. The combined direct anti-tumor activity and indirect effects on host metabolism are likely to contribute to the observed, enhanced anti-tumor effects in this DIO setting. We provide evidence that SDX-7320 stimulates weight loss and improves obesity-driven metabolic dysfunction, corroborating previous non-clinical and clinical reports with MetAP2 inhibitors (18,20). The combination of multi-model tumor biology, transcriptomics and metabolomics represents a significant strength. The head-to-head comparison with tirzepatide is a strength, as it reveals the role of host metabolic factors on tumor growth, highlights a unique mechanism of action of SDX-7320, and reinforces the clinical and translational relevance of the work. Potential bias was mitigated by the use of independent Contract Research Organizations to execute the experiments using industry-standard endpoints. However, it also is important to acknowledge certain limitations of the work. For example, we do not yet have evidence that the SDX-7320-mediated changes in the expression of immune system-related genes revealed by RNA-sequencing are functionally important in this setting. Future experiments are envisioned to explore the role of alterations to the tumor immune microenvironment and systemic immune system following administration of SDX-7320. Moreover, with the exception of the EO771 model of TNBC, all experiments were carried out in male mice or rats. We cannot exclude sex-based differences, particularly for immune and metabolic outcomes.

Given the widespread prevalence of baseline and treatment-induced metabolic dysfunction in cancer patients and its reported deleterious impact on the progression of many types of solid tumors, combining standard-of-care oncology therapy with a complementary agent that has both anti-tumor and metabolic correcting effects warrants consideration for clinical development. The fact that SDX-7320 demonstrated inhibition of obesity-accelerated tumor growth in multiple mouse models highlights the promise of using an agent that has both direct anti-tumor effects and beneficial effects on host metabolism. Currently, SDX-7320 (evexomostat) is being clinically investigated as a first-in-class metabo-oncology agent for patients with metastatic TNBC in combination with chemotherapy and in patients with metastatic hormone receptor-positive, HER-2-negative breast cancer in combination with standard-of-care targeted therapies (NCT05570253 and NCT05455619, respectively).

## Acknowledgements

The authors would like to acknowledge Agilux Laboratories (Worcester, MA) for bioanalytical support, Gwosdow Associates Science Consultants for help with editing the manuscript, SBH Sciences (Natick, MA) for biomarker measurements, The Jackson Lab (Bar Harbor, ME), Neosome Life Sciences (Billerica, MA), and Rincon Bio (Lehi, UT) for in vivo studies, PanomeBio (St. Louis, MO) and both Watershed AI, LLC and Rancho Biosciences for bioinformatics support.

## Funding

This work was funded by SynDevRx, Inc.

## Notes

**Conflict of Interest Disclosure Statement:** P. Cornelius reports patents 10646463, 11273142, and 11612577 issued; patents for 18/036,565, PCT/US2023/064550, PCT/US2023/072023, and PCT/US2023/080080 pending; and P. Cornelius is an employee of SynDevRx, Inc. and holds stock options in SynDevRx, Inc. B.A. Mayes reports patents for PCT/US2023/064550 and PCT/US2023/080080 pending, and B.A. Mayes is an employee of SynDevRx, Inc. and owns stock options in SynDevRx. J.S. Petersen reports patents 9173956, 9433600, 9750737, 9757373, 10010544, 10588904, 11304944, 9320805, 9585909, 9730955, 9895449, 10159692, 10722532, 9969722, and 10287277 issued. P.J. Dufour is an employee of SynDevRx, Inc., with ownership and options on equity. A.J. Dannenberg reports personal fees from SynDevRx during the study and ownership of stock options in SynDevRx. J.M. Shanahan reports patents 9173956, 9433600, 9750737, 9757373, 10010544, 10588904, 11304944, 10646463, and 11273142, respectively, and J.M. Shanahan is a full-time employee of SynDevRx and holds both stock and stock options. B.J. Carver reports patents for 18/036,565 and PCT/US2023/064550 pending, and B.J. Carver is a full-time employee of SynDevRx, Inc. and holds both stock and stock options. No disclosures have been reported by the other authors.

### Competing Interest Statement

P. Cornelius reports patents 10646463, 11273142, and 11612577 issued; patents for 18/036,565, PCT/US2023/064550, PCT/US2023/072023, and PCT/US2023/080080 pending; and P. Cornelius is an employee of SynDevRx, Inc. and holds stock options in SynDevRx, Inc. B.A. Mayes reports patents for PCT/US2023/064550 and PCT/US2023/080080 pending, and B.A. Mayes is an employee of SynDevRx, Inc. and owns stock options in SynDevRx. J.S. Petersen reports patents 9173956, 9433600, 9750737, 9757373, 10010544, 10588904, 11304944, 9320805, 9585909, 9730955, 9895449, 10159692, 10722532, 9969722, and 10287277 issued. P.J. Dufour is an employee of SynDevRx, Inc., with ownership and options on equity. A.J. Dannenberg reports personal fees from SynDevRx during the study and ownership of stock options in SynDevRx. J.M. Shanahan reports patents 9173956, 9433600, 9750737, 9757373, 10010544, 10588904, 11304944, 10646463, and 11273142, respectively, and J.M. Shanahan is a full-time employee of SynDevRx and holds both stock and stock options. B.J. Carver reports patents for 18/036,565 and PCT/US2023/064550 pending, and B.J. Carver is a full-time employee of SynDevRx, Inc. and holds both stock and stock options. No disclosures have been reported by the other authors.

