## Supplementary Figures for "Evexomostat (SDX-7320), a Methionine Aminopeptidase Type 2 (METAP2) Inhibitor, Stimulates Weight Loss and Inhibits Obesity-Accelerated Tumor Growth"

**Supplementary Figure S1. Anti-obesity and anti-diabetes activity of SDX-7320 in DIO mice.** Male C57Bl/6J mice (individually housed) were fed a high-fat/high-sucrose diet for 12 weeks (Harlan Diet TD.06414; beginning at 6 weeks of age) then randomized based on body weight (n=6/group). SDX-7320 or vehicle (5% mannitol in water) were administered subcutaneously (Q4D) for a total of 8 doses. Body weight was measured every other day, and food intake per cage was measured on a weekly basis. Intraperitoneal glucose tolerance test (IPGTT) was conducted 24 days after the first dose of SDX-7320, using 1 g/kg of glucose and blood glucose measurements were made using a hand-held glucometer (blood samples were obtained by tail nick). The glucose area under the curve (AUC) was calculated using GraphPad, version 10.0. Adipose tissue depots as well as liver were dissected and weighed at the end of the study. Plasma was prepared from blood samples collected at the end of the study and sent to IDEXX for measurement of ALT, cholesterol and triglycerides. Data were analyzed using repeated measures two-way ANOVA with Geisser-Greenhouse correction and Sidak's multiple comparisons for results in Panels A, B and D whereas Kruskal-Wallis one-way ANOVA with Dunn's multiple comparisons test was used to analyze results in Panels C, and E – I (GraphPad, v10.0; adjusted p values are reported as follows: *p < 0.05, **p < 0.01, ***p < 0.005 versus vehicle-treated mice).

(A) (B)

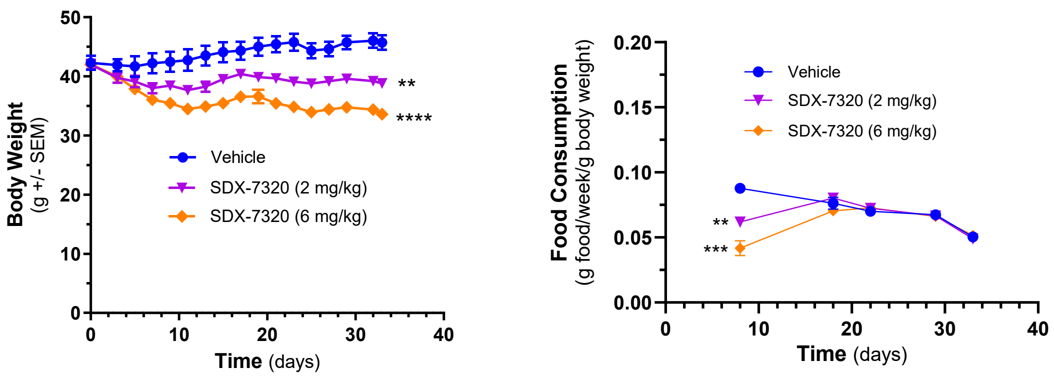

(C)

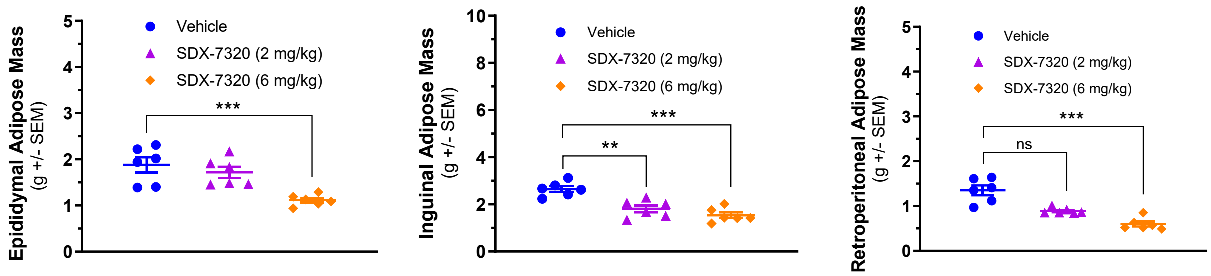

Epididymal Adipose Mass
(g +/- SEM)
Vehicle
SDX-7320 (2 mg/kg)
SDX-7320 (6 mg/kg)
***
Inguinal Adipose Mass
(g +/- SEM)
**
***
Retroperitoneal Adipose Mass
(g +/- SEM)
ns
***


(D)

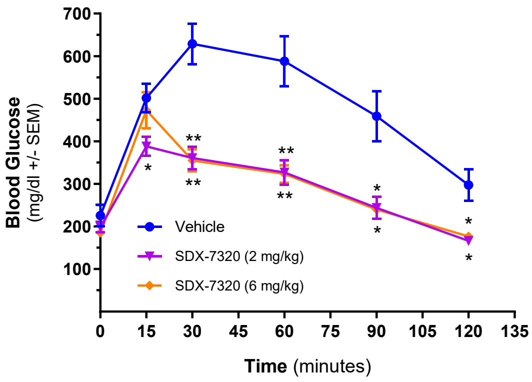

Blood Glucose
(mg/dl +/- SEM)
Vehicle
SDX-7320 (2 mg/kg)
SDX-7320 (6 mg/kg)
Time (minutes)
0 15 30 45 60 75 90 105 120 135


(E)

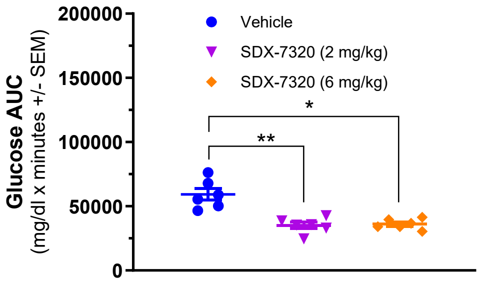

Glucose AUC
(mg/dl x minutes +/- SEM)
Vehicle
SDX-7320 (2 mg/kg)
SDX-7320 (6 mg/kg)
**
*


(F)

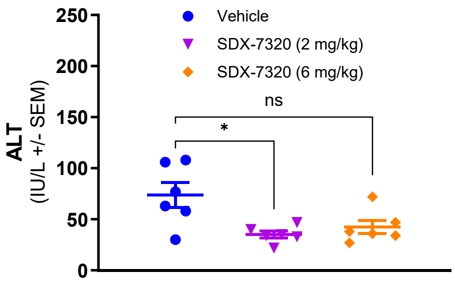

ALT
(IU/L +/- SEM)
Vehicle
SDX-7320 (2 mg/kg)
SDX-7320 (6 mg/kg)
*
ns


(G)

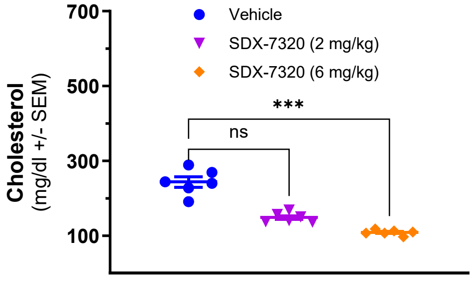

Cholesterol
(mg/dl +/- SEM)
Vehicle
SDX-7320 (2 mg/kg)
SDX-7320 (6 mg/kg)
ns
***


(H)

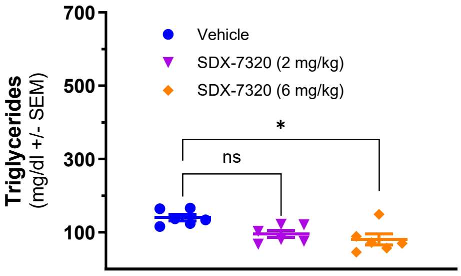

Triglycerides
(mg/dl +/- SEM)
Vehicle
SDX-7320 (2 mg/kg)
SDX-7320 (6 mg/kg)
ns
*


(I)

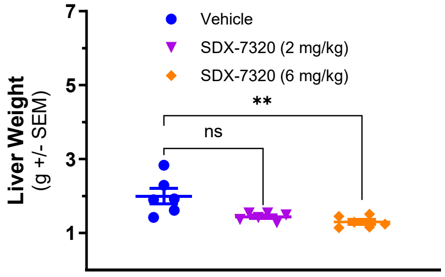

Liver Weight
(g +/- SEM)
Vehicle
SDX-7320 (2 mg/kg)
SDX-7320 (6 mg/kg)
ns
**

**Supplementary Figure S2**. **Effect of SDX-7320 on insulin sensitivity in DIO mice.**

Diet-induced obese mice (male C57Bl/6j, n=6/group) were dosed subcutaneously with vehicle (5% mannitol in water) or with SDX-7320. Four days after dosing, food was removed from the mouse cages prior to an insulin-tolerance test (ITT). Humulin-R was diluted in 0.9% NaCl and delivered intra-peritoneally at 7.5 mL/kg at t=0, to achieve 0.75 U/kg. Blood glucose was measured with an EasyMax V glucometer at t=0 and then 15-, 30-, 45-, 60-, and 90 minutes post-insulin (Panel A). ITT results were normalized to baseline glucose and re-analyzed (Panel B). Data were analyzed using repeated measures two-way ANOVA with Geisser-Greenhouse correction and Sidak's multiple comparisons (GraphPad, v10.0; *$p < 0.05$, **$p < 0.01$versus vehicle-treated mice).

(A) (B)

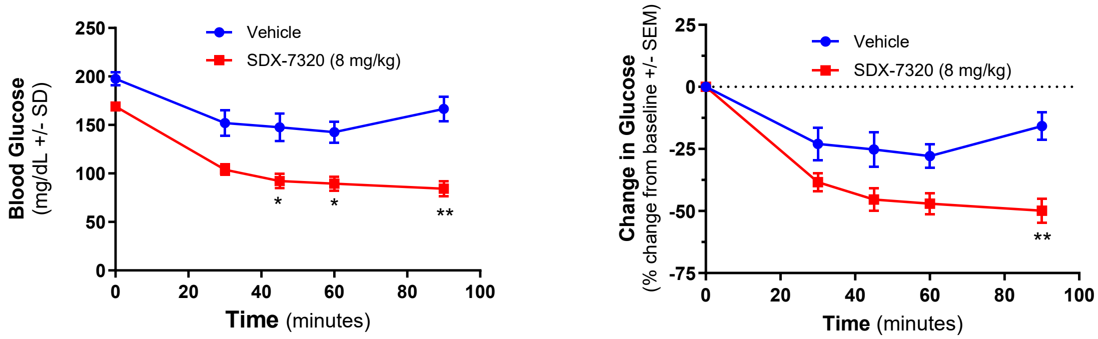

**Supplementary Figure S3. Anti-obesity activity of SDX-7320 and SDX-7539 in obese rats.** Obese male Sprague-Dawley DS (Levin) rats maintained on a high-fat/high-sucrose diet were dosed by subcutaneous injection with vehicle (PBS), SDX-7320 (0.3, 1.0, 3.0 mg/kg) or SDX-7539 (1.0, 3.0 mg/kg) once every 4 days (N=3/group). Food intake was measured weekly for each animal (Panel A). After 12 doses (day 48) rats were fasted for four hours, then subjected to an oral glucose tolerance test (OGTT) using 2 g/kg glucose. Blood samples taken during the OGTT were analyzed for glucose and insulin levels at the indicated time points, as well as calculation of the AUC for each endpoint (Panels B-E). Adipose tissue depots (epididymal, inguinal and retroperitoneal) were dissected and weighed at the end of the study (day 68; Panels F-H). Time-dependent data in panels A, B and D were analyzed using repeated measures two-way ANOVA with Geisser-Greenhouse correction and Sidak's multiple comparisons test whereas Kruskal-Wallis one-way ANOVA with Dunn's multiple comparisons test was used for analysis of data in panels C, E - H (GraphPad, v10.0; *$p < 0.05$, **$p < 0.01$ versus vehicle-treated mice).

(A)

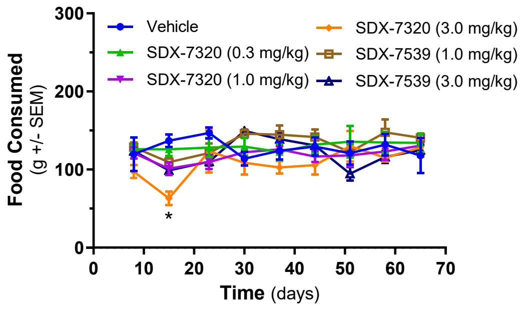


(B)

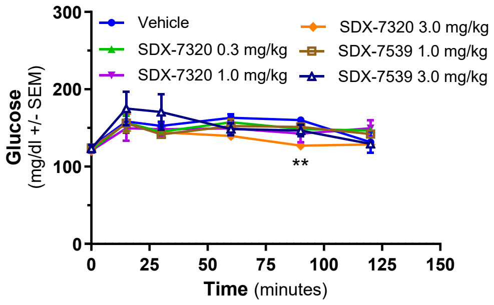


(C)

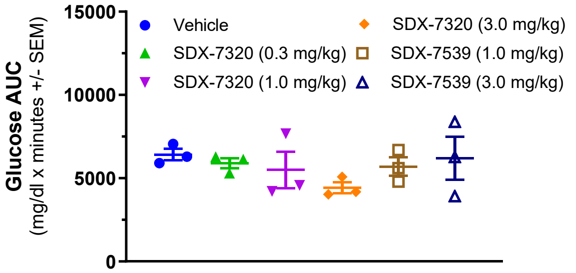


(D)

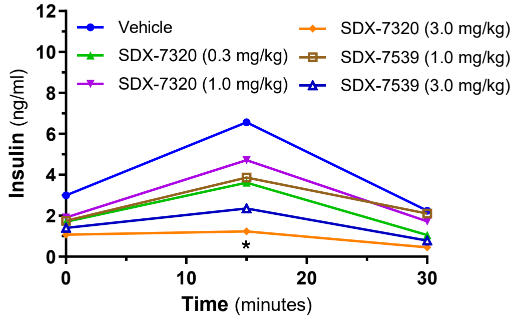

(E)

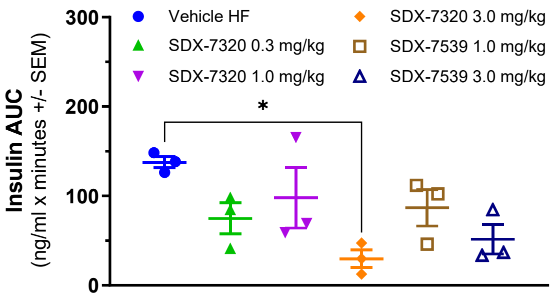
Insulin AUC
(ng/ml x minutes +/- SEM)
300
200
100
0
Vehicle HF
SDX-7320 0.3 mg/kg
SDX-7320 1.0 mg/kg
SDX-7320 3.0 mg/kg
SDX-7539 1.0 mg/kg
SDX-7539 3.0 mg/kg
*

(F)

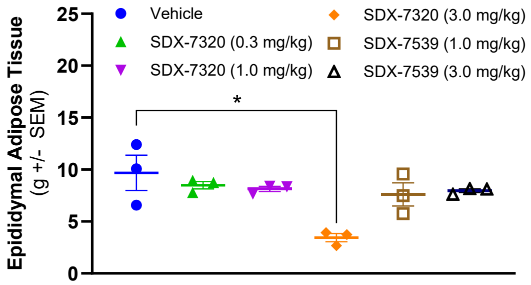
Epididymal Adipose Tissue
(g +/- SEM)
25
20
15
10
5
0
Vehicle
SDX-7320 (0.3 mg/kg)
SDX-7320 (1.0 mg/kg)
SDX-7320 (3.0 mg/kg)
SDX-7539 (1.0 mg/kg)
SDX-7539 (3.0 mg/kg)
*

(G)

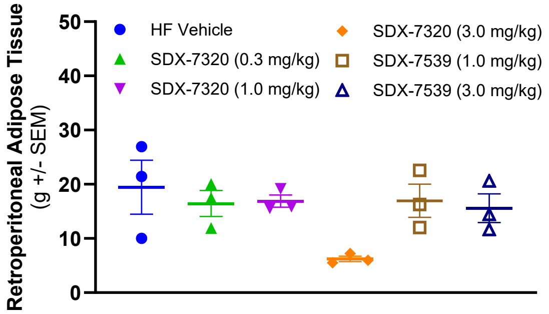
Retroperitoneal Adipose Tissue
(g +/- SEM)
50
40
30
20
10
0
HF Vehicle
SDX-7320 (0.3 mg/kg)
SDX-7320 (1.0 mg/kg)
SDX-7320 (3.0 mg/kg)
SDX-7539 (1.0 mg/kg)
SDX-7539 (3.0 mg/kg)

(H)

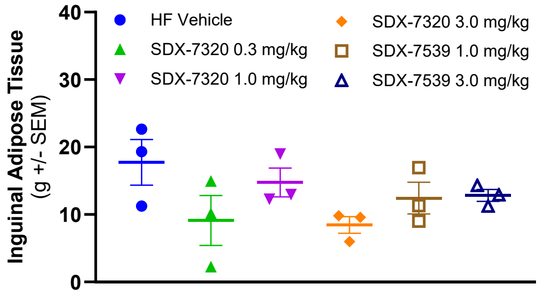
Inguinal Adipose Tissue
(g +/- SEM)
40
30
20
10
0
HF Vehicle
SDX-7320 0.3 mg/kg
SDX-7320 1.0 mg/kg
SDX-7320 3.0 mg/kg
SDX-7539 1.0 mg/kg
SDX-7539 3.0 mg/kg

**Supplementary Figure S4. Effect of SDX-7320 on EO771 mammary gland tumor growth in lean and obese mice.** Diet-induced obese mice (female C57Bl/6, surgically ovariectomized, n=10 group) or their lean littermates (n=9/group) were injected with EO771 cells (a model of triple-negative breast cancer) into the fourth mammary gland ($1x10^5$ cells/mouse; 50/50 Matrigel/PBS). When tumors were approximately 60 $mm^3$, mice were dosed subcutaneously with vehicle (5% mannitol in water) or with SDX-7320 (8 mg/kg, Q4D dissolved in vehicle). Body weight and tumor volume were measured twice/week (Panels A-E). Two weeks after dosing was initiated, mice were euthanized, and adipose tissue was dissected and weighed (Panels F-H). Time-dependent data in panels A - E were analyzed using repeated measures two-way ANOVA with Geisser-Greenhouse correction and Sidak's multiple comparisons test whereas Mann-Whitney non-parametric test was used for analysis of data in panels F – H (GraphPad, v10.0; *$p < 0.05$, **$p < 0.01$ versus vehicle-treated mice).

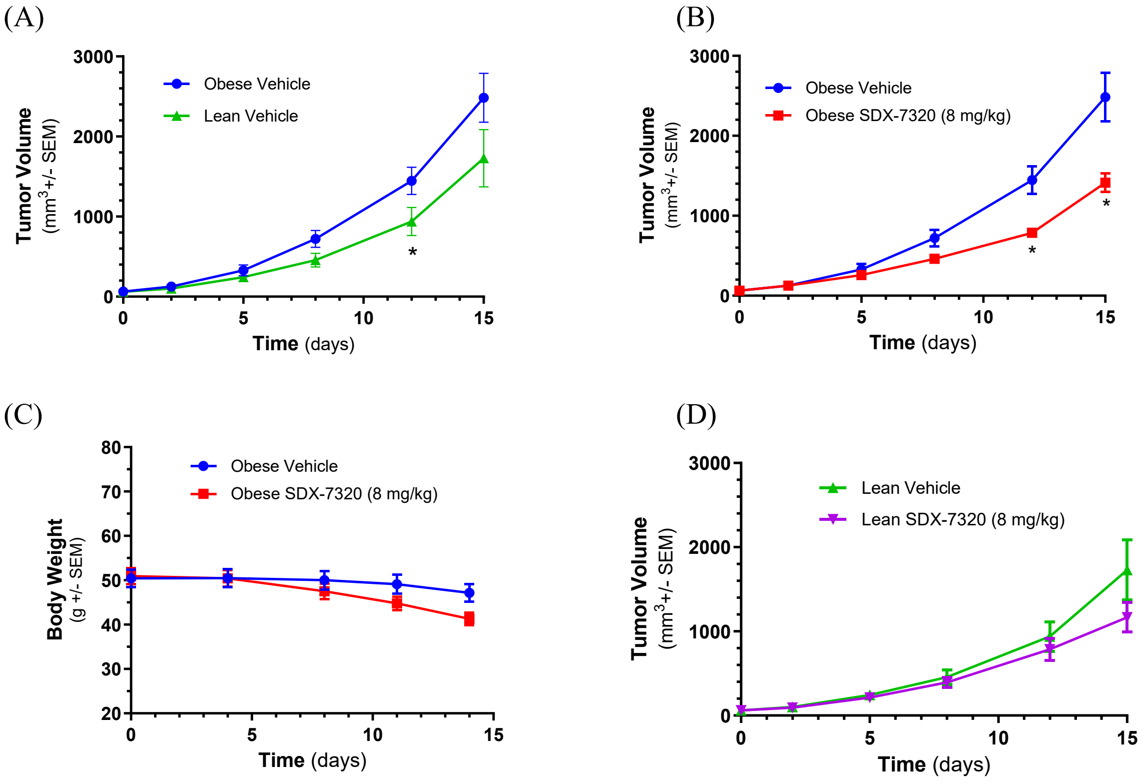

(E)

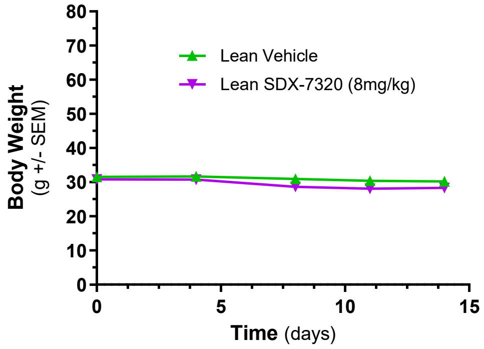
Body Weight
(g +/- SEM)
80
70
60
50
40
30
20
10
0
0
5
10
15
Time (days)
Lean Vehicle
Lean SDX-7320 (8mg/kg)

(F)

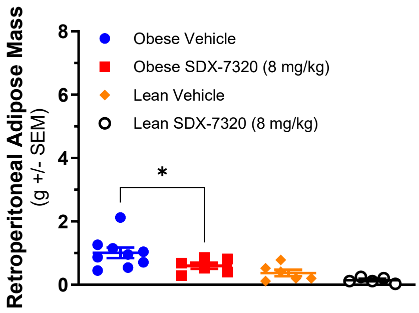
Retroperitoneal Adipose Mass
(g +/- SEM)
8
6
4
2
0
*
Obese Vehicle
Obese SDX-7320 (8 mg/kg)
Lean Vehicle
Lean SDX-7320 (8 mg/kg)

(G)

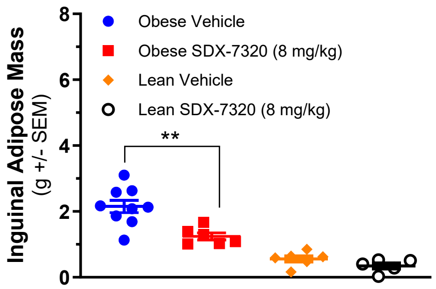
Inguinal Adipose Mass
(g +/- SEM)
8
6
4
2
0
**
Obese Vehicle
Obese SDX-7320 (8 mg/kg)
Lean Vehicle
Lean SDX-7320 (8 mg/kg)

(H)

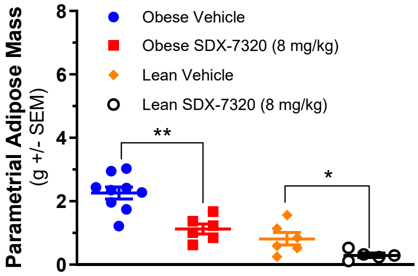
Parametrial Adipose Mass
(g +/- SEM)
8
6
4
2
0
**
*
Obese Vehicle
Obese SDX-7320 (8 mg/kg)
Lean Vehicle
Lean SDX-7320 (8 mg/kg)

**Supplementary Figure S5. Effect of SDX-7320 vs. vehicle on gene expression in MC38 tumors from obese vs. lean mice.** Diet-induced obese mice (male C57Bl/6) or their lean littermates were injected in the right rear flank with MC38 tumor cells (2 x $10^5$ cells/mouse). When tumors were approximately 100 $mm^3$, mice were dosed subcutaneously Q4D with vehicle (5% mannitol in water) or with SDX-7320 (6 mg/kg). Two weeks after dosing was initiated, mice were euthanized, and tumors were dissected and placed into RNALater. Bulk RNA-Seq was conducted on tumor samples (N=4 per group) and data was analyzed using DESeq2. Panels A and B, volcano plots for SDX-7320 vs. vehicle for obese and lean mice, respectively.

(A)

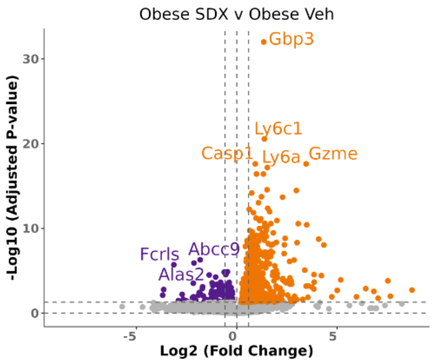


(B)

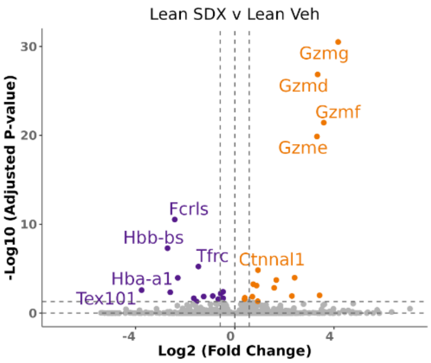

**Supplementary Figure S6. Changes in plasma metabolites from SDX-7320- and tirzepatide-treated obese mice bearing MC38 tumors**. This Venn diagram shows the number of significantly altered metabolites observed exclusively in tirzepatide (TZP)-treated mice vs vehicle (red), SDX-7320-treated mice vs vehicle (blue green), or those that overlapped between comparisons.

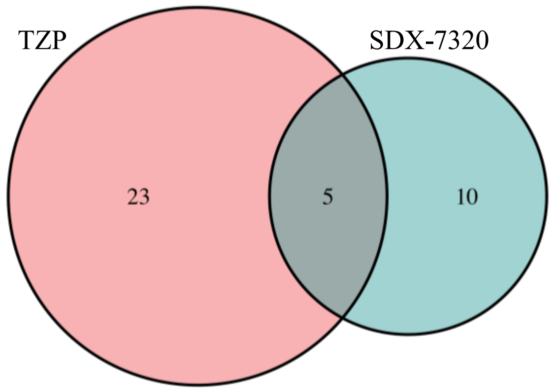

**Supplementary Table S1. Pharmacokinetic parameters for SDX-7539 in obese Levin rats.** At 1-, 2-, 4- and 24-hours post-dose, blood samples were collected, and plasma was prepared for analysis of SDX-7539 levels using LC/MS (N=3/group). Pharmacokinetic parameters $C_{max}$ and AUC were calculated using GraphPad Prism v10. Statistical analysis was carried out using Brown-Forsythe and Welch's one-way ANOVA with Dunnett's T3 multiple comparisons (GraphPad Prism v10).

| **SDX-7320 Dose** mg/kg, (N) | **SDX-7539 $C_{max}$** ng/ml +/- SD | **SDX-7539 AUC** ng*h/ml +/- SD |
|---|---|---|
| 0.3 (3) | 0.32 +/- 0.10 | 4.62 +/- 0.97 |
| 1.0 (3) | 1.86 +/- 1.33 | 25.5 +/- 13.4 |
| 3.0 (3) | 2.73 +/- 0.26 | 39.9 +/- 2.72 |
| **SDX-7539 Dose** mg/kg, (N) | | |
| 1.0 (3) | 76.2 +/- 8.82 | 168 +/- 11.6 |
| 3.0 (3) | 230 +/- 19.0 | 1048 +/- 425 |

| **Statistical Analysis of SDX-7539 AUC Data** | | | | |
|---|---|---|---|---|
| **Dunnett's T3 multiple comparisons test** | **Mean Diff.** | **95% CI of diff.** | **Summary** | **Adjusted P Value** |
| SDX-7320 (0.3 mg/kg) vs. SDX-7320 (1.0 mg/kg) | -20.92 | -87.16 to 45.33 | ns | 0.3825 |
| SDX-7320 (0.3 mg/kg) vs. SDX-7320 (3.0 mg/kg) | -34.46 | -41.91 to -27.00 | *** | 0.0006 |
| SDX-7320 (0.3 mg/kg) vs. SDX-7539 (1.0 mg/kg) | -163.8 | -221.5 to -106.0 | ** | 0.0064 |
| SDX-7320 (0.3 mg/kg) vs. SDX-7539 (3.0 mg/kg) | -1043 | -3147 to 1060 | ns | 0.1842 |
| SDX-7320 (1.0 mg/kg) vs. SDX-7320 (3.0 mg/kg) | -13.54 | -80.35 to 53.27 | ns | 0.6579 |
| SDX-7320 (1.0 mg/kg) vs. SDX-7539 (1.0 mg/kg) | -142.8 | -192.1 to -93.60 | *** | 0.0010 |
| SDX-7320 (1.0 mg/kg) vs. SDX-7539 (3.0 mg/kg) | -1023 | -3127 to 1082 | ns | 0.1910 |
| SDX-7320 (3.0 mg/kg) vs. SDX-7539 (1.0 mg/kg) | -129.3 | -187.7 to -70.89 | * | 0.0105 |
| SDX-7320 (3.0 mg/kg) vs. SDX-7539 (3.0 mg/kg) | -1009 | -3113 to 1095 | ns | 0.1953 |
| SDX-7539 (1.0 mg/kg) vs. SDX-7539 (3.0 mg/kg) | -879.7 | -2984 to 1225 | ns | 0.2467 |

| Statistical Analysis of SDX-7539 $C_{max}$ Data | | | | |
|---|---|---|---|---|
| **Dunnett's T3 multiple comparisons test** | **Mean Diff.** | **95% CI of diff.** | **Summary** | **Adjusted P Value** |
| SDX-7320 (0.3 mg/kg) vs. SDX-7320 (1.0 mg/kg) | -1.540 | -8.141 to 5.061 | ns | 0.5673 |
| SDX-7320 (0.3 mg/kg) vs. SDX-7320 (3.0 mg/kg) | -2.410 | -3.345 to -1.475 | ** | 0.0033 |
| SDX-7320 (0.3 mg/kg) vs. SDX-7539 (1.0 mg/kg) | -75.88 | -119.5 to -32.22 | * | 0.0169 |
| SDX-7320 (0.3 mg/kg) vs. SDX-7539 (3.0 mg/kg) | -229.7 | -323.7 to -135.6 | ** | 0.0086 |
| SDX-7320 (1.0 mg/kg) vs. SDX-7320 (3.0 mg/kg) | -0.8700 | -7.577 to 5.837 | ns | 0.8921 |
| SDX-7320 (1.0 mg/kg) vs. SDX-7539 (1.0 mg/kg) | -74.34 | -118.5 to -30.19 | * | 0.0180 |
| SDX-7320 (1.0 mg/kg) vs. SDX-7539 (3.0 mg/kg) | -228.1 | -322.4 to -133.9 | ** | 0.0088 |
| SDX-7320 (3.0 mg/kg) vs. SDX-7539 (1.0 mg/kg) | -73.47 | -117.1 to -29.80 | * | 0.0181 |
| SDX-7320 (3.0 mg/kg) vs. SDX-7539 (3.0 mg/kg) | -227.3 | -321.3 to -133.2 | ** | 0.0088 |
| SDX-7539 (1.0 mg/kg) vs. SDX-7539 (3.0 mg/kg) | -153.8 | -224.1 to -83.50 | ** | 0.0053 |

**Supplementary Table S2. Differentially expressed genes in MC38 tumors from lean and obese mice (see Excel file).** Diet-induced obese mice (male C57Bl/6) or their lean littermates were injected in the right rear flank with MC38 tumor cells. When tumors were approximately 100 $mm^3$, mice were dosed subcutaneously with vehicle or with SDX-7320 (6 mg/kg). Approximately, two weeks after dosing was initiated, mice were euthanized and tumors were dissected and placed into RNALater. Bulk RNA-Seq was conducted on tumor samples (N=4 per group). Comparisons of lean SDX-7320 vs. vehicle and obese SDX-7320 vs. vehicle are shown in separate tabs. Cells shaded orange indicate genes whose expression increased following treatment with SDX-7320, that overlap between lean and obese mice. Cells shaded purple indicate genes whose expression decreased following treatment with SDX-7320, that overlap between lean and obese mice.

**Supplementary Table S3. Effect of SDX-7320 and tirzepatide on levels of metabolites in plasma (see Excel file).** Diet-induced obese mice (male C57Bl/6j) were injected in the right rear flank with MC38 tumor cells. When tumors were approximately 60 $mm^3$, mice were dosed subcutaneously with vehicle, SDX-7320 (6 mg/kg, Q4D) or tirzepatide (30 nmol/kg, QD). Eighteen days after initiating dosing, animals were euthanized, and plasma was prepared (N=7-8/group) for untargeted metabolomics analysis. Select metabolites from SDX-7320 or tirzepatide treatment groups were significantly different from the vehicle group. Metabolites that increased are shown in red; molecules that were reduced are shown in blue.
